# Plant miRNAs Anti-*Staphylococcus aureus:* Therapeutic Perspective

**DOI:** 10.1101/2024.06.13.598803

**Authors:** Charfeddine Gharsallah

## Abstract

*Staphylococcus aureus* is a pathogen that has developed resistance to each new antibiotic introduced for half a century especially through the acquisition of the *mecA* gene. This bacterial resistance to antibiotics represents a major public health problem. New revolutions are underway, in particular the design of drugs and vaccines targeting the system for detecting the regulatory quorum of the accessory gene (*agr*). It has been shown that the pathogenicity and resistance of *S. aureus* can be modulated through the intervention of this system. For this reason, we propose in this present work, a new therapeutic design based on an *in silico* study to identify plant miRNAs that could target this system as well as the *mecA* gene based on recent studies showing the inter-realm regulation of human transcripts by plant miRNAs. Out of a total of 20643 miRNAs from mature plants, we identified 29 miRNAs, obtained by the selection criteria MFE ≥ -25 kcal / mol, which could potentially target selected genes of *S. aureus*. Fifteen of them were selected on the basis of their thermodynamic stability. Interestingly, The seeds of ptc - miR171g and ptc - miR171h (*Populus trichocarpa*) both of which almost similar mature miRNA sequence was found to target UCCC region of *RNAIII* gene. miRNAs from plants targeting *S. aureus* have the potential to be developed as an alternative therapy of the future. To our knowledge, it is the first therapeutic alternative of the future via the *in silico* identification of plant miRNAs targeting pathogenic bacteria.

**Graphical Abstract:** 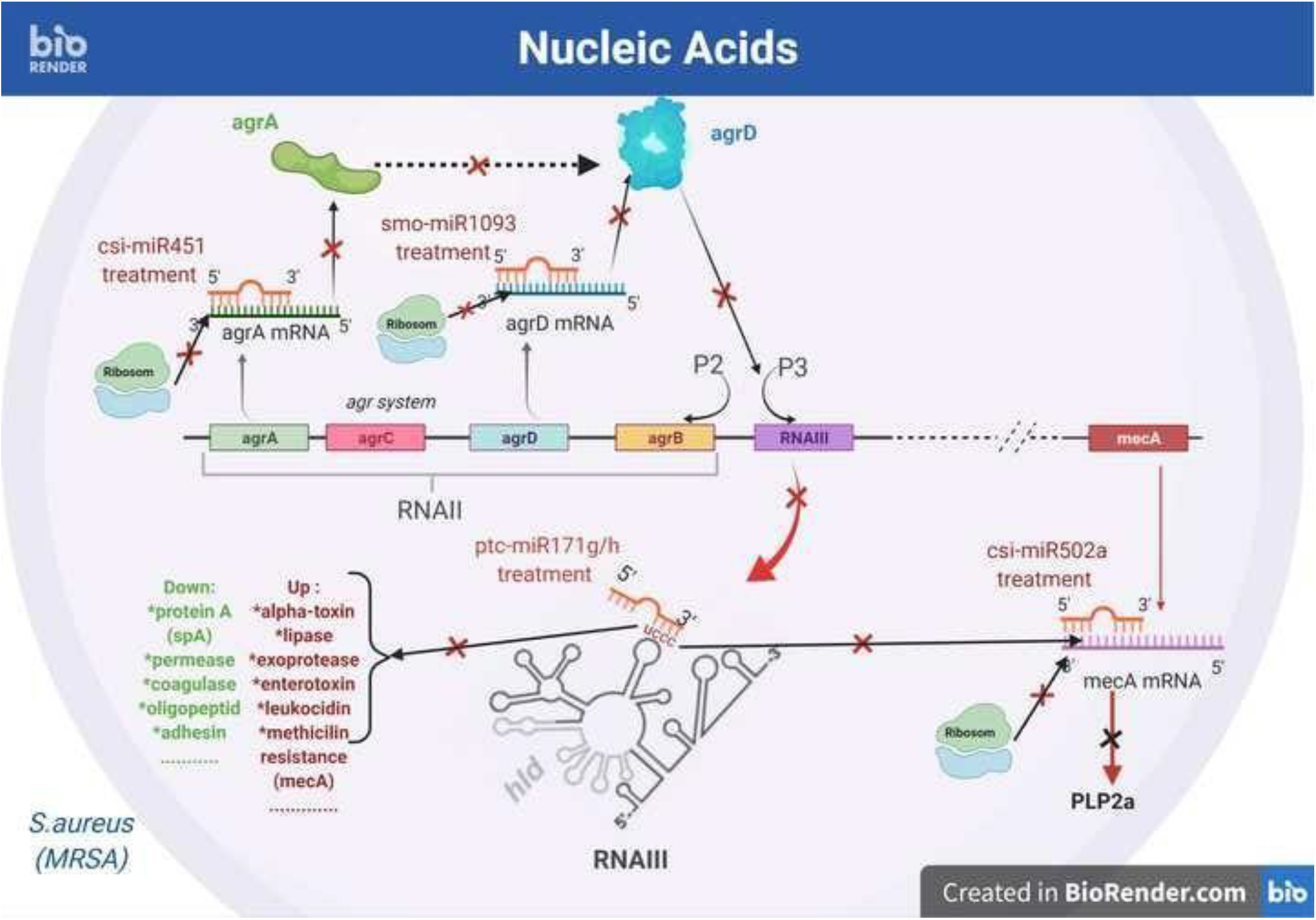

## 1. Introduction

*Staphylococcus aureus* (*S. aureus*) is a very versatile pathogen, able to inhabit the skin and mucous membranes like a commensal (Novick, 2003). It can also invade the bloodstream and different tissues, causing various serious diseases (Krismer and Peschel, 2011). *S. aureus* is the main cause of nosocomial and community infections in the world (Mandal et al., 2015), with conditions ranging from skin invasion to fatal systemic illnesses, such as pneumonia and endocarditis (Thammavongsa et al., 2015). Several studies have attributed such wide range of infections to its vast arsenal of virulence factors (adhesins, toxins and enzymes) (Li et al., 2014; Tuchscherr and Löffler, 2015). These factors are under the control of the quorum-sensing accessory gene regulator (*agr*) system (Li et al., 2014).

This *agr* system intervenes during the post-transcriptional regulation of virulence factors and biofilm formation but also during the heterogeneous resistance of *S. aureus* to methicillin (MRSA) (Kavanaugh and Horswill, 2016; Mohsenzadeh et al., 2015; Singh and Ray, 2014). The *agr* operon is based on two different promoters, P2 and P3, and generates respectively two primary transcripts, (i) RNAII coding for *agrB, agrD, agrC* and *agrA*) and (ii) a post-transcriptional regulator called *RNAIII* (Tan et al., 2018). The *RNAIII* regulates the expression of several transcripts among them alpha-hemolysin (*hla*), beta-hemolysin (*hlb*), toxic shock syndrome toxin-1 (*tsst*), and staphylococcal toxin serine protease (*sspA*). The *S. aureus*’ *RNAIII* is present in a single copy and has an UCCC consensus sequence, that initiate its hybridization to the mRNA. Through various structural domains, *RNAIII* acts either as an activator or as a repressor of mRNA targets. For example, studies carried out by Queck et al. 2008 suggested that *RNAIII* positively regulates the expression of *mecA* gene, whereas it negatively regulates *spA* gene (Huntzinger et al., 2005). Accordingly, *agr* system is considered as an interesting therapeutic target (Tan et al., 2018). and an effective method for weakening the virulence and/or resistance of staphylococcal pathogens.

Several attempts have been made to produce effector molecules of the *agr* system, Gov et al. 2001 synthesized the RNA-III inhibitory peptide (RIP and native RIP) which effectively suppresses diseases caused by *S. aureus* by inhibiting the transcription of *RNAII* and *RNAIII*. Indeed, RIP reduces bacterial adhesion to cells and prevents biofilm formation (Gov et al., 2001).

MiRNAs are known to regulate a wide variety of functions during cellular development and also patho-physiological processes (O’Brien et al., 2018). They are known to interfere, via mRNA silencing, with protein translation through base pairing to the 3’-untranslated regions (3’-UTR) or the coding region, leading to the cleavage or the downregulation of the corresponding gene (Sanchita et al., 2018).

There has been an increasing focus on miRNAs as a therapeutic tool for different disease treatment strategies (Parsi et al., 2015). In this context, Teng et al., 2018 have shown that gma-miR396e is involved in the modulation of the composition of the intestinal microbiota and their metabolites, which can be used to inhibit mouse colitis. Likewise, ath-miR-167a (*Arabidosis thaliana*) was found to be regulating the expression of *SpA*C from *Lactobacillus rhamnosus*, which allows it to remain on the surface of the mucosa and hence contributes to intestinal recovery. In addition, mdo-miR-7267-3p down regulates *ycnE* expression, increasing the production of indole-3-carboxaldehyde (IA3) and leading to reduced intestinal permeability. It has been reported that exosomes from bovine modulate the proportion of cecum microbes by stimulating the growth of certain bacteria such as *Tenericutes* and *Firmicutes* in mice (Zhou et al. 2019).

We performed herein an *in silico* study to identify the binding sites of putative plant miRNAs to the *agr* system (*agrA, agrD, RNAIII*) as well as the *mecA* gene, as a new therapeutic route to control the virulence / resistance of *S. aureus* strains.

## 2. Materials and Methods

### 2.1. *In silico* identification and analysis of plants miRNA potentially targeting *agr* system (*agrA, agrD* and *RNAIII*) and *mecA* gene

#### 2.1.1. *S. aureus* genome

The methicillin-resistant *S. aureus* genome sequence was downloaded from the NCBI database (Refseq ID: HE579067.1). It corresponds to the complete sequence of the genome of Swiss hospital origin (Switzerland). This sequence was used as a reference against plant miRNAs targeting the regions of different *agr* system and *mecA* genes. We have chosen from all the *agr* system genes the regulatory gene *RNAIII* as well as the *agrA* (cytoplasmic regulator) and *agrD* (propeptide) genes; the latter being strongly involved in the activation of *RNAIII* (Gilot et al., 2002).

#### 2.1.2. Predicted plant miRNAs

Targeted miRNAs used in this work were found in the psRNATarget (Dai et al., 2018) and MepmiRDB (medicinal plant) (Yu et al., 2019) databases as well as other publications (Khan et al. 2020; Lin et al. 2018; Kalarikkal & Sundaram 2021). All of these targets were anticipated by the RNAhybrid (Krü and Rehmsmeier, 2006) to predict potential plant miRNA binding sites of the *mecA, RNAIII, agrA* and *agrD* genes. Default parameters have been used to calculate minimum free energies “MFEs≤ -25 kcal/mol” to ensure that the binding of miRNA to target mRNA can supersede the local secondary structures (Kertesz et al., 2007; Mückstein et al., 2006). The heliX constraint was adjusted in seed sequences between 2 and 8 bases from the 5 ‘end of miRNA. Non-canonical G-U base pairing was allowed since it is well tolerated in the deletion of target mRNA mediated by miRNA (Saxena et al., 2003). Conservation of miRNA binding sites across *S. aureus* strains are used to assess the specificity of miRNA-target interactions. Hence, we aligned the miRNA binding site sequence within the *S. aureus* genome (Refseq ID: HE579067.1) with global strains of *S. aureus* using MEGA software, version 10.1 (Stecher et al., 2020).

#### 2.1.3. Associating network

The network was generated by Cytoscape v3.0.2 (Cline et al., 2007) to set up associations between plant miRNAs identified via RNAhybrid (Krü and Rehmsmeier, 2006) with their potential binding sites in regions of the *agr* system and *mecA* gene. The algorithm MCODE28 from the Cytoscape Plugin Cluster Maker allowed the network partitioning into layers, where the most discriminating miRNAs with corresponding genes and plants were linked. This discrimination is based on the secondary structure predictions by MFOLD web server (Zuker, 2003), and the thermodynamic stability at 37°C of miRNAs was calculated using RNAfold (Hofacker, 2003).

#### 2.1.4. The UCCC consensus sequence of *RNAIII* and ptc-miR171g/h Tertiary Structure Prediction and Docking

The UCCC consensus sequence of *RNAIII* and ptc - miR171g / h tertiary structures have been generated using RNAComposer tools (Biesiada et al., 2016), and subsequently docked using HNADOCK server (He et al., 2019).

## 3. Results

### 3.1. Plant microRNAs: exploitation of a regulatory mechanism between kingdoms to target the genome of *S. aureus* (*mecA, agrA, agrD* and *RNAIII*)

#### 3.1.1. Plant miRNA contain *S. aureus* targeting miRNAs

The entire process of miRNA selection, mature miRNA sequence retrieval, target prediction, specificity of miRNA-target interaction and secondary structure analysis is depicted as a flow chart in Figure 1.

**Figure 1.**
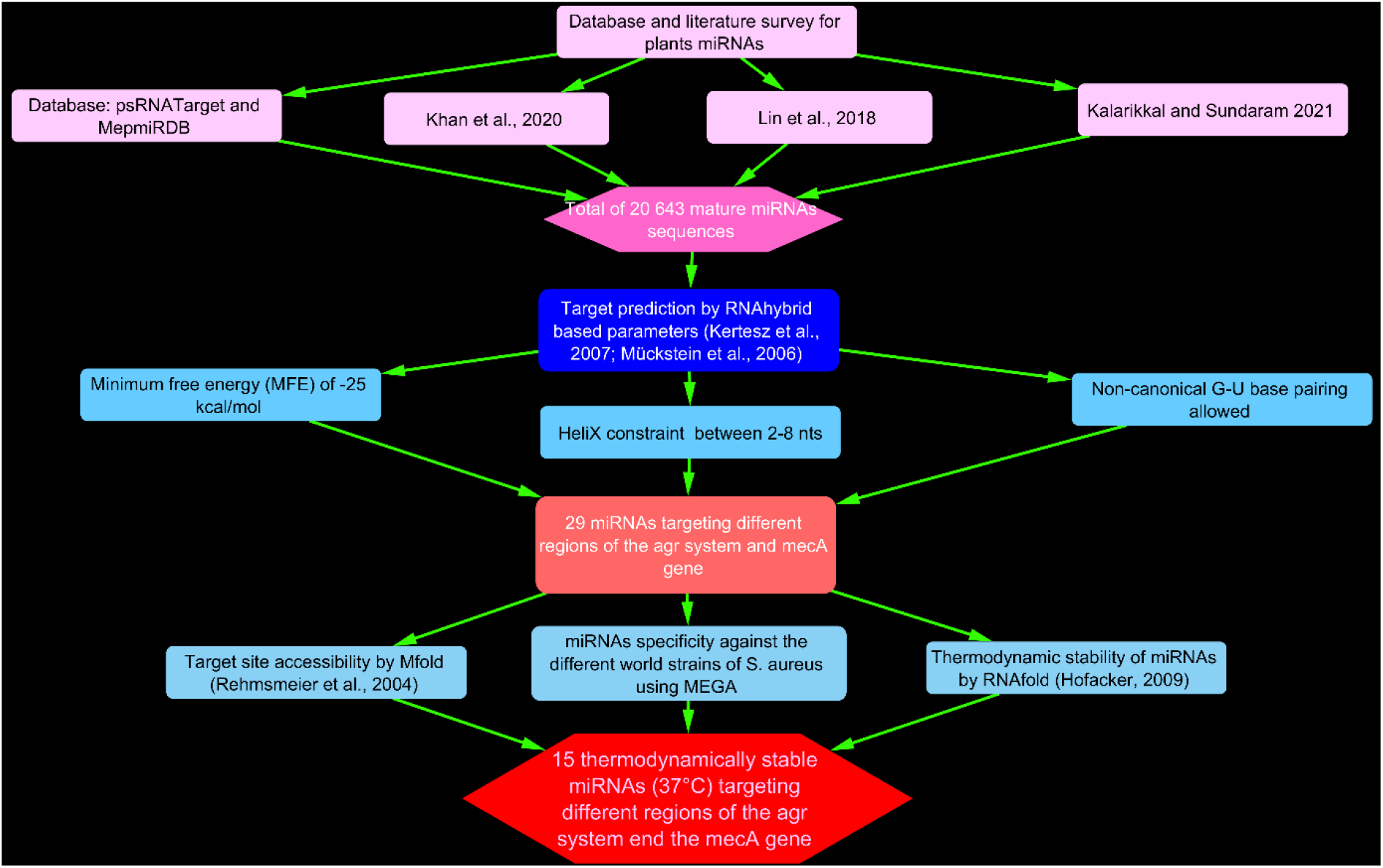
Flowchart summarizing the *in silico* approach used for the prediction of plant miRNAs and the analysis of their targets

A total of 20,643 mature miRNAs **[seeSupporting information_ SI]** were taken as input for this study and subjected to target site prediction against *RNAIII, agrA, agrD* and *mecA* gene sequences with RNAhybrid software. Using rigorous selection criteria (Figure 1), we first obtained 29 plant miRNAs that could target the genes of *S. aureus*. These miRNAs, their origin and their targets on *S. aureus* genome are represented on the “plant-miRNAs-*S. aureus* genes” multilayer network (Figure2). We have identified plant miRNAs of edible origin (potato, rice, soy, orange, pomegranate …) as well as of medicinal origin (ginkgo, medicago, dendranthema …). Interestingly, the vast majority of identified plant miRNAs (14 out of 29 miRNAs) had target sites within *mecA* gene, and 7 out of 29 had target sites within agrA et *RNAIII* genes respectively. The *agrD* gene was only targeted by a single miRNA (smo-miR-1093). Out of 29 miRNAs, 5 were identified in *Citrus sinensis* corresponding to csi-miRN502a, csi-miR3953, csi-miRN807 and csi-miRN263 targeting *mecA* gene and csi-miRN451 targeting *agrA* gene. Three mtr-miR5244, mtr-miR5554 and mtr-miR399t identified in *Medicago truncatula*, were able to target gene encoding *agrA*. Besides, three ptc-miR171g, ptc-miR171h and ptc-miR394 identified in *Populus trichocarpa*, were able to target different genes encoding *RNAIII*I (miR171g/h), and *mecA* gene respectively. ath-miR775 targeting *agrA* gene and ath-miR8181 targeting *mecA* gene were identified in *Arabidopsis thaliana*. Similarly, two miRNAs were selected for *Selaginella moellendorffii*. These miRNAs target genomic regions of genes encoding *agrA* (smo-miR1093) and *mecA* (smo-miR1097). The other species have only one miRNA targeting the *S. aureus* genes (Figure2).

All plants identified belong to different taxonomic families **[see Supporting information_Table S1]**. A second selection was made considering both the structures obtained by the selection criteria MFE = <25 kcal / mol by the RNAhybrid software and the thermodynamic stability (37°C) of the miRNAs obtained by the RNAfold software (described in methods) (**see Supporting information_Table S2**). Fifteen miRNAs were selected on the basis of their thermodynamic stability which are shown in the Figure 2 (branches in red color) and Table 1. Thus, among the 14 miRNAs targeting the mecA gene, 7 were retained on the basis of their thermodynamic stability (bra-miR6032-3p, csi-miR263-3p, csi-miR502a-3p, dmo-miR40-5p, lyc-miR30 -3p, smo-miR1097 and aly-miR166f-5p). Among the 7 miRNAs targeting the agrA gene, 4 were retained on the basis of their thermodynamic stability (mtr-miR5244, mtr-miR5554a/b/c-3p, csi-miR451-5p and smi-miR112-3p). Among the 7 miRNAs targeting the RNAIII gene, 3 were retained on the basis of their thermodynamic stability (ptc-miR171g-5p, ptc-miR171h-5p and gma-miR1524). smo-miR-1093 targeting the agrD gene was selected based on its thermodynamic stability. The secondary structures of miRNA-target mRNA duplex for all 15 miRNAs along with target site sequences are shown in figure 3.

**Table 1.**
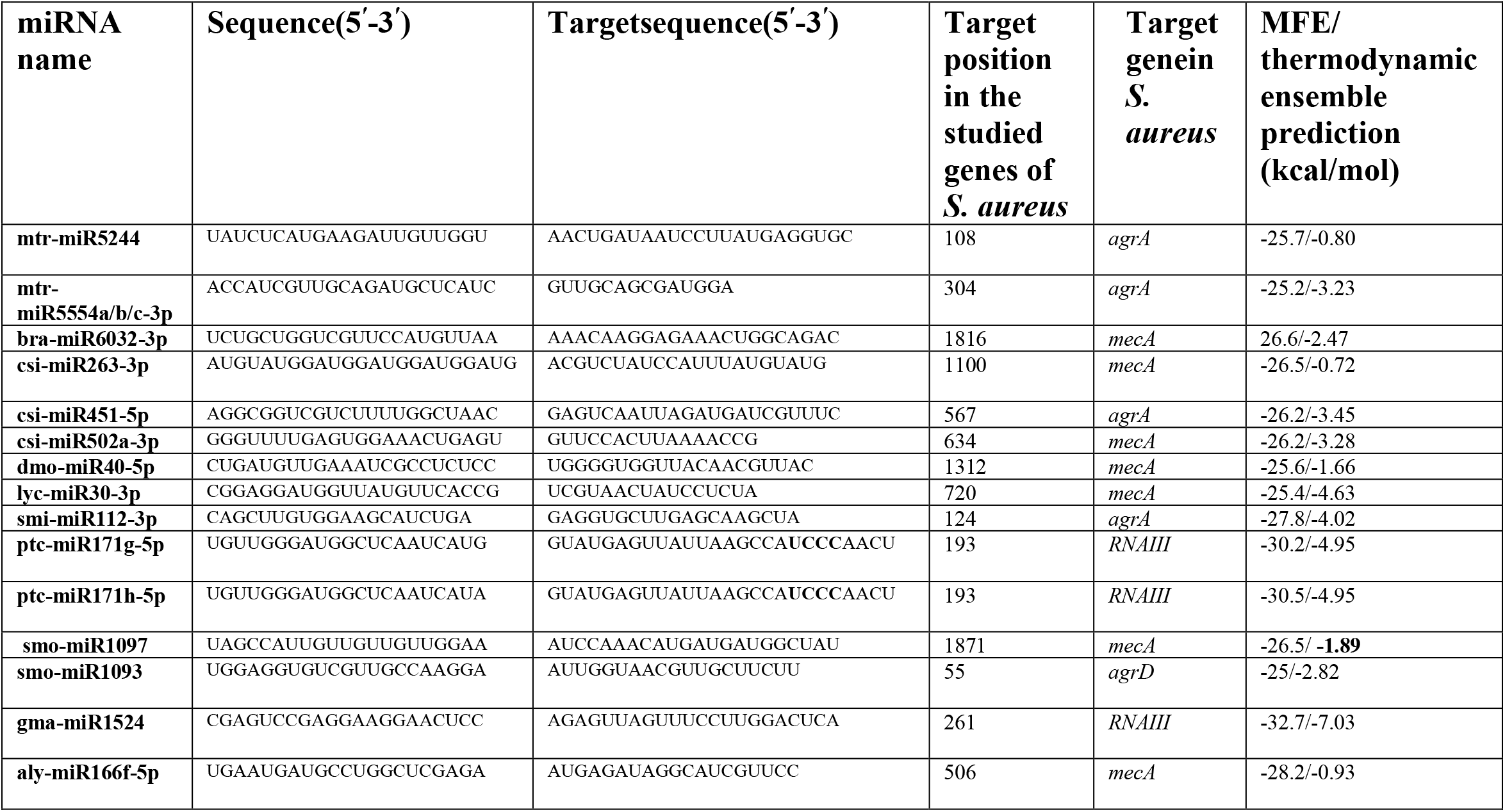
List of plant miRNAs targeting *RNAIII, agrA, agrD* and *mecA* gene of *S aureus* genome.

**Figure 2.**
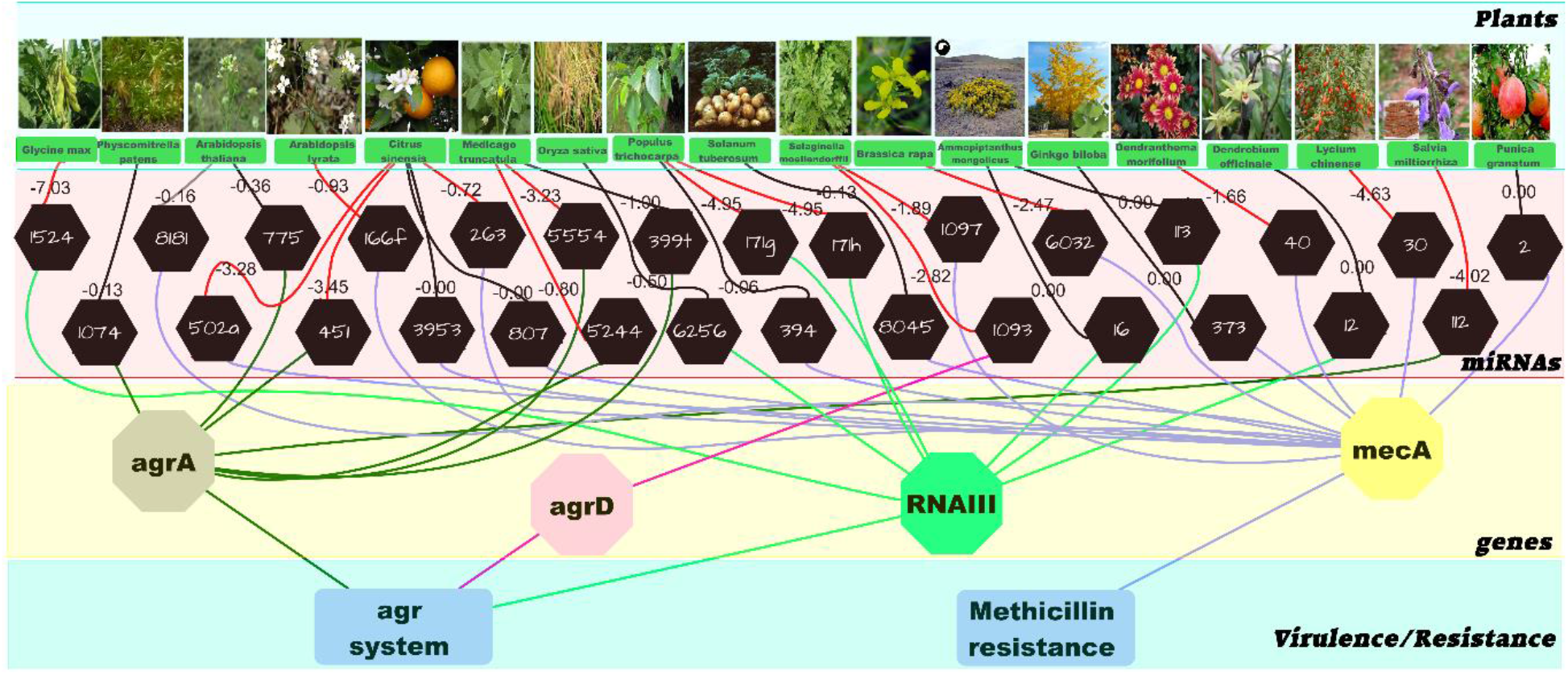
“Predicted plant miRNAs-*S. aureus* genes” association network. Black hexagon nodes represent plant miRNAs. Pink, yellow, green and grey hexagons represent *S. aureus* target genes. The values of thermodynamic stability are represented at the black (unstable) and red (stable) branches. The clear green branches connect the predicted plant miRNAs with the *RNAIII* gene; the dark green branches connect the predicted plant miRNAs with the *agrA* gene; the pink branches connect the predicted plant miRNAs with the *agrD* gene and the purple branches connect the predicted plant miRNAs with the *mecA* gene.

**Figure 3.**
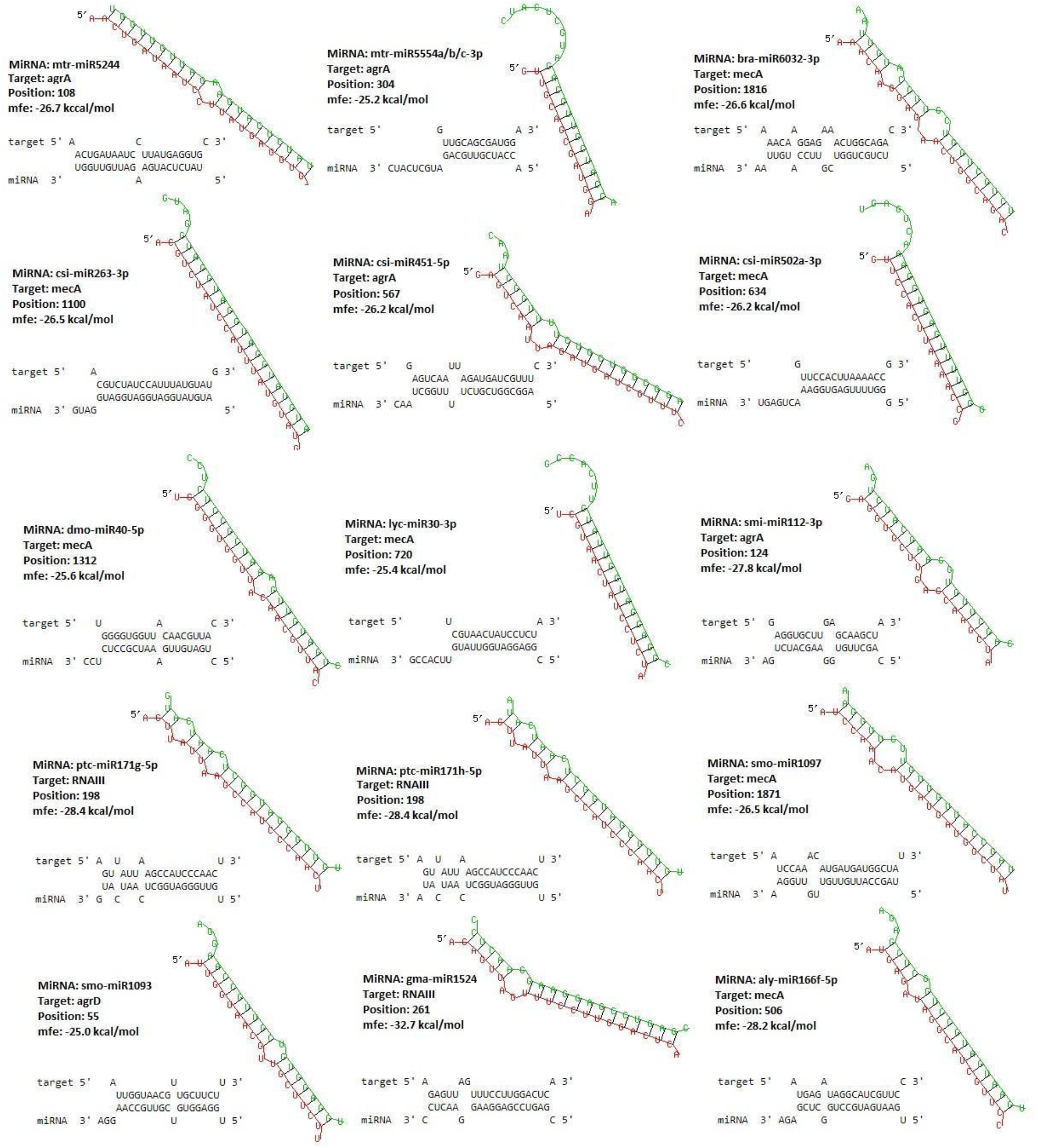
Schematic depiction of stable RNA hybrid structures formed between 15 plant miRNAs (in green) with target sites in *S. aureus* genes (in red). The relative nucleotide position of target sites for each miRNA in *S. aureus* genome is shown left the alignments.

#### 3.1.2 ptc - miR171g / h targets the consensus recognition site “UCCC” of *RNAIII*

As previously mentioned, *S. aureus RNAIII* is present in a single copy on the genome and has the UCCC consensus sequence, found at 7, 13 and 14 and recognized to initiate the *RNAIII* / mRNA interaction. This RNA positively regulates the expression of the resistance gene *mecA* (Huntzinger et al., 2005). Disruption of ribosomal fixation event at the recognized site will lead to the premature termination of translation, thereby inhibiting the synthesis of *mecA*, and therefore resistance to methicillin. Interestingly, the seeds of ptc - miR171g and ptc miR171h both of which almost similar mature miRNA sequence was found to target UCCC region of *RNAIII* gene (Table 2). Modeling of the bidirectional and three-dimensional structure showed that the 5 ‘end of miR-ptc - miR171g / h established complementary contacts with *RNAIII* (UCCC) with an MFE = -223.54 kcal / mol which exceeds the MFE of the *RNAIII* bond “UCCC”-*mecA* (-205.95 kcal / mol) (Figure 4). The UCCC site is conserved in *Staphylococcaceae* and mutations in this region lead to abrogation of target regulation (Desgranges et al., 2019; Geissmann et al., 2009; Liu et al., 2011). Despite these findings, we assumed a mutational event affecting each nucleotide of the UCCC site in order to see its impact on the ptc-miR171g/h-*RNAIII* interaction. The results reveal that the mutational event for each position of the UCCC site does not affect the efficiency of the ptc-miR171g/h-*RNAIII* interaction (Figure 5). Modeling of the three-dimensional structure showed that ptc-miR171g/h maintains the establishment of complementary contacts with RNAIII in the presence of the VCCC, UDCC, UCDC and UCCD mutations with values MFEs = -209.92; -242.66; -217.23 and -211.38 kcal/mol respectively (Figure 5). These values also exceed the MFE of the *RNAIII*-mecA bond (-205.95 kcal/mol).

**Figure 4.**
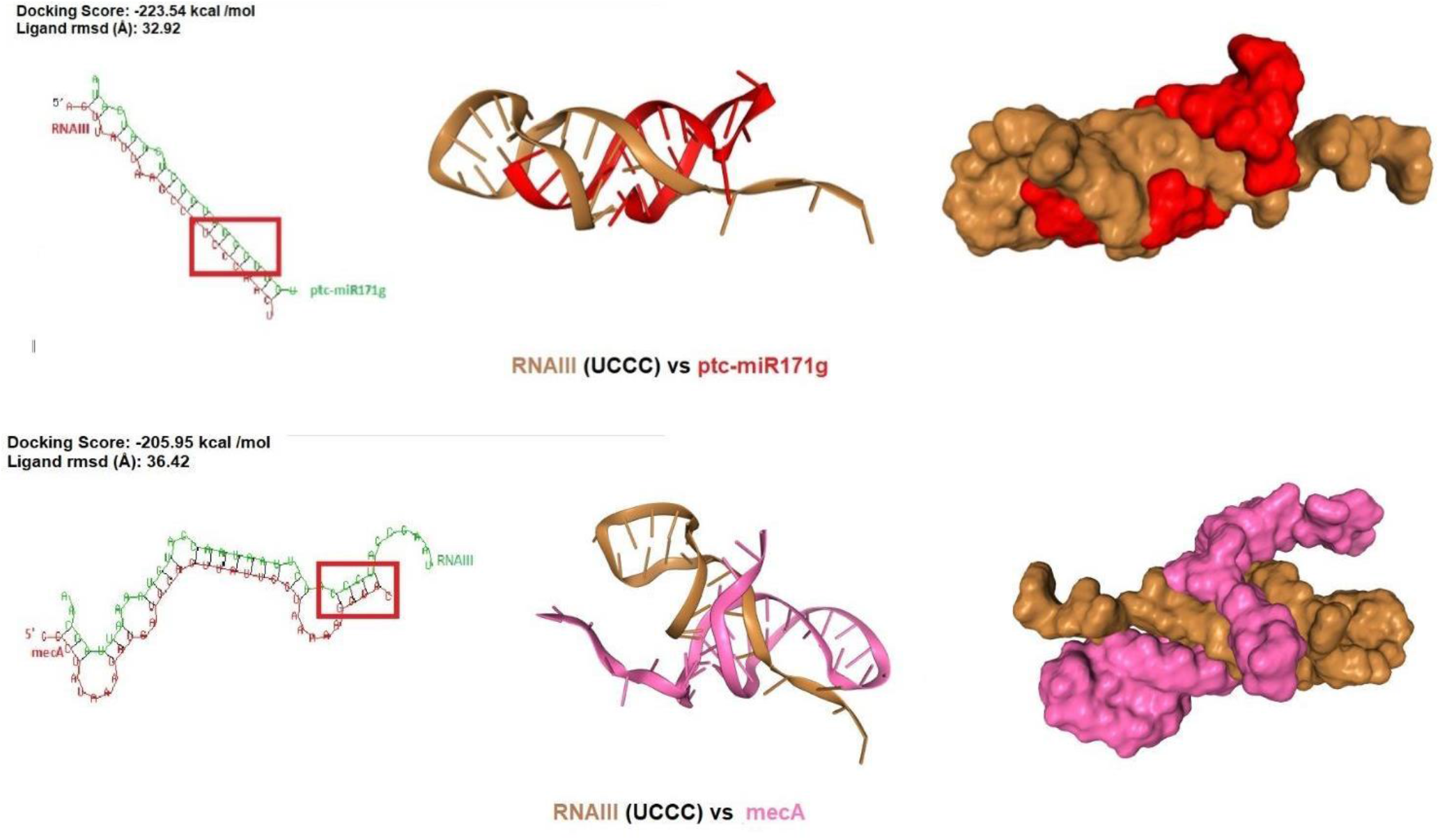
The binding site for ptc-miR171g/h hybridizes with the UCCC recognition site of *RNAIII* gene. Low MFE (-223.54 kcal / mol) should promote the binding of ptc-miR171g / h with the UCCC recognition site and therefore block all binding of *RNAIII* with the genes it regulates (*mecA, spA*, etc.).

**Figure 5.**
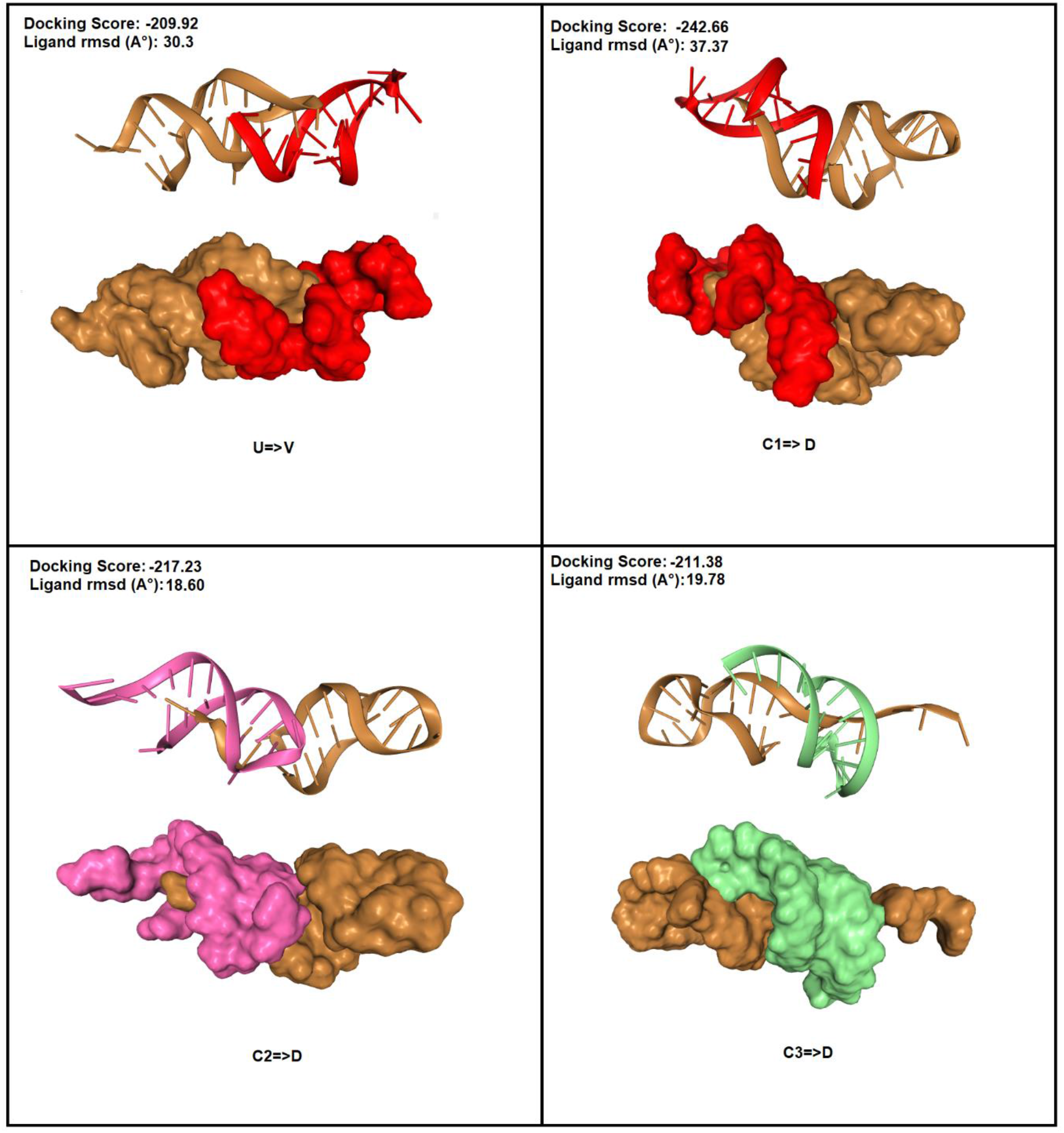
ptc-miR171g/h hybridizes with RNAIII in the presence of mutations (V not T and D not C) on the UCCC recognition site of the RNAIII gene. A low MFE should promote the binding of ptc-miR171g/h with RNAIII and thus allow the disruption of post-transcriptional regulation with the genes it regulates (mecA, spA, alpha-toxin etc.).

#### 3.1.3 Target specificity of plant miRNAs to different strains isolated from all over the world

To investigate the specificity of the selected miRNAs against the genes studied in this work in different strains of *S. aureus*, we aligned the miRNA target sequences identified using the strain (HE579067.1) with sequences of *S. aureus* isolated from other countries such as USA, South Africa, Germany, Korea and China. It has been observed that global strains of *S. aureus* share 100% sequence similarity with the genes studied in the Swiss strain (HE579067.1). Due to this, all the 15 miRNA target sites were highly conserved across all isolates of S. *aureus* (Figure 6). Based on this, seven miRNAs target the mec A gene, 4 target miRNAs the *agrA* gene, 2 target miRNAs the *RNAIII* gene and only one miRNA target the *agrD* gene (Figure 6). We can suggest that these different miRNAs are able to suppress methicillin resistance and virulence of *S. aureus* and can be considered as future drugs thus replacing the different antibiotics.

**Figure 6.**
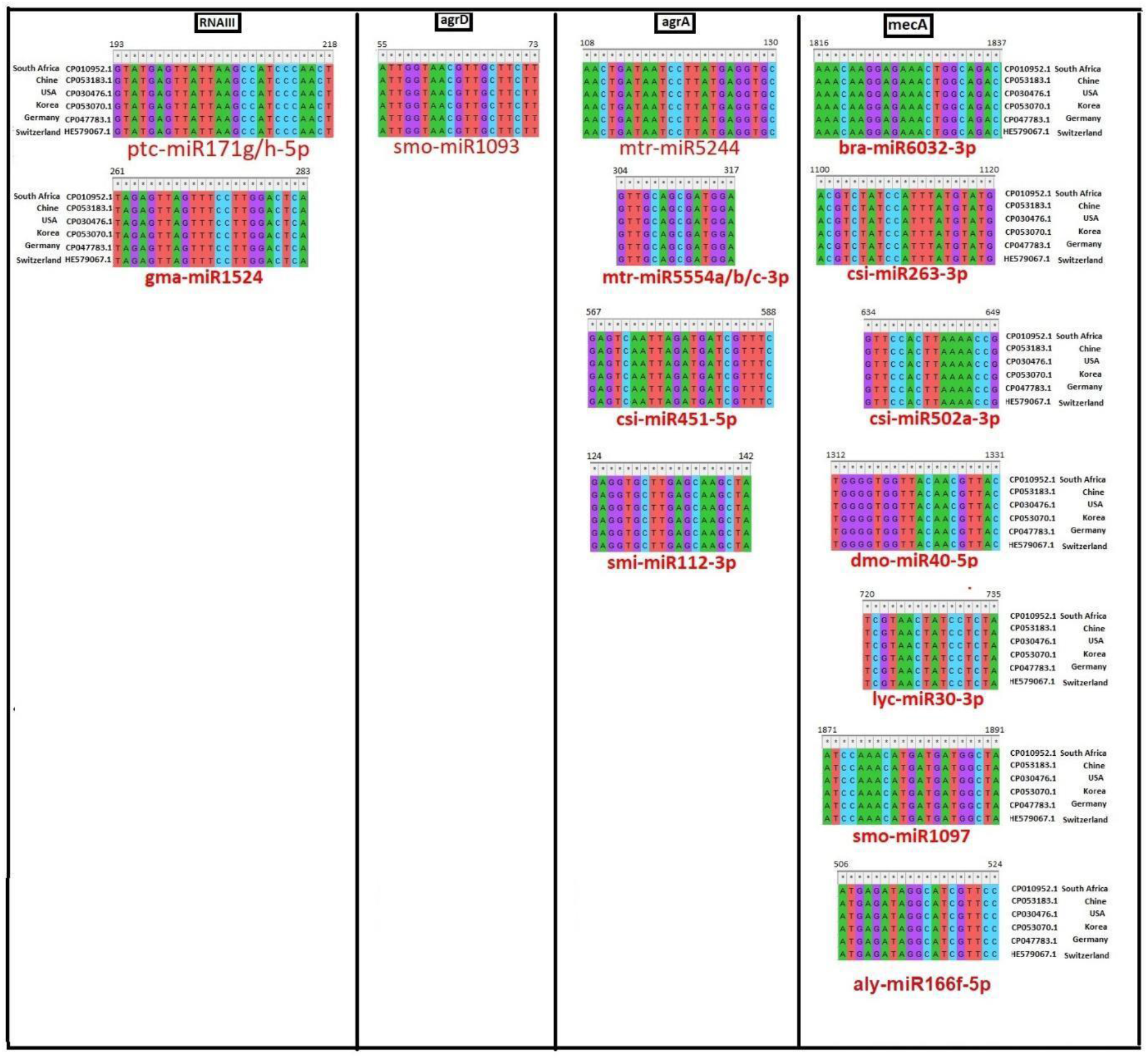
Segregation of plant miRNAs based on specificity against *S. aureus* genes. Noting well that the seed sequences are located at the 3′ end of the mRNA sequence where 5′ end of miRNA binds.

#### 3.1.4. Target site accessibility of plant miRNA binding sites

The accessibility of the target site to miRNA binding is one of the critical keys determining successful miRNA mediated target mRNA suppression (Long et al., 2007; Witkos et al., 2011). It has been experimentally validated that the local secondary structure between 17 nt upstream and 13 nt downstream of the miRNA binding site can influence target site accessibility to miRNAs (Kertesz et al., 2007). Hence, we assessed the local secondary structure formation at the miRNA target site along with 17 nt upstream and 13 nt down-stream sequences from *S. aureus* genes using MFOLD webserver. Out of the 15 plant miRNA binding sites in *S. aureus*, we found 3 miRNA target sequences having at least one predicted secondary structure with MFE -9 kcal/mol **[SeeSupporting information_Table S3]**. The target sites of smi-miR112-3p and mtr-miR5244 overlapped with local secondary structures with MFE of -4.7 kcal/mol and -10.4 kcal/mol, respectively. Since all the 15 miRNA binding sites within *S. aureus* genome formed secondary structures with MFE ≤ 25 kcal/mol, we anticipate that plant miRNAs could supersede the local secondary structures to allow their own binding.

## 4. Discussion

Despite their small size, miRNA molecules have enormous potential due to their ability to regulate the expression of target genes. It has been shown that miRNA profiles are correlated with many human diseases and pathological stages, which attracted more attention from scientists and the pharmaceutical industry. The ability of plant miRNAs to regulate gene expression in a cross-kingdom mode has already been demonstrated for different diseases (Díez-Sainz et al., 2021; Mangukia et al., 2021; Teng et al., 2018). Zhang et al. (2012) were the first to demonstrate these intriguing properties of miRNA for plant molecules. They suggested that rice miR168a is a potential target for low density lipoprotein receptor 1 (LDLRAP1) adapter protein mRNA, and that feeding mice rice has shown upregulation of this receptor expression in the blood and liver. A recent study suggested that transcription factor 7 (TCF7) is involved in the Wnt signaling pathway and is overexpressed in breast cancer (Chin et al., 2016). This study showed that the abundance of miR159 from *Arabidopsis thaliana* and soybean (Glycine max) is inversely correlated with the progression of cancer cells. Recently, the localization of miR-156c and miR-159a has been demonstrated in edible nanoparticles derived from nuts and those miRNAs involved in the mammalian TNF-α signaling pathway in adipocytes and regulate inflammation (Aquilano et al., 2019). Regarding studies on antiviral therapy via plant miRNAs, Zhou et al. (2015; 2020) showed that feeding the boiled honeysuckle decoction causes an elevation of the level of MIR2911 in the serum and lung of mice infected with the SARS-CoV-2 virus as well as influenza A (IAV) and therefore inhibits their replication and reduce mortality (Zhou et al., 2020, 2015). Recently, studies have shown that diet modulates the composition and function of the gut microbiota in humans (Teng et al., 2018). Indeed, they identified gma-miR396e involved in the modulation of the composition of the intestinal microbiota and their metabolites and inhibit mouse colitis. This miRNA promotes the growth of *Lactobacillus rhamnosus* (LGG) at least in part by inhibiting the expression of LexA (gene encoding the protein that represses number of genes involved in the response to DNA damage: response SOS). Likewise, ath-miR-167a (*Arabidosis thaliana*) regulates the expression of *SpA*C from *Lactobacillus rhamnosus*, which allows it to remain on the surface of the mucosa. In addition, mdo-miR-7267-3p downregulates ycnE expression, increasing the production of indole-3-carboxaldehyde (IA3) and leading to reduced intestinal permeability. Similar findings have been discussed by Díez-Sainz et al., 2021, that a diet would be a main source of absorption of xeno-miRNAs (xenomiRs) in different organs. All studies have allowed us to think that miRNAs of plant origin can be used as future therapy against pathogenic bacteria mainly nosocomial, given their implication in the regulation of bacterial genes in our body. In this work, we further explore the possibility of exploiting the regulation of the cross kingdom by plant miRNAs to treat *S. aureus* infections. This strategy is incriminated by the framework of minimizing the misuse of antibiotics and therefore the proliferation of multi-resistant strains. In this line, we have identified plant miRNAs that could potentially target key genes of *S. aureus*. We first extracted these datasets to identify plant miRNAs sequenced to date. Next, using a target prediction pipeline with strict selection criteria, we identified 15 miRNAs, based on their thermodynamic stability, that bind to the genes selected in this work. Interestingly, ptc miR171g and ptc - miR171h, derived from *Populus trichocarpa*, which almost similar mature miRNA sequence was found to target the UCCC region of the *RNAIII* gene of the *S. aureus agr* system upregulating *mecA* resistance gene expression (Queck et al., 2008). We have also shown that mutations that can affect this site do not affect the hybridization of ptc - miR171g/h with RNAIII. It should be noted that this tree was commonly used by many Indian tribes in North America who particularly appreciated it for its antiseptic, anti-inflammatory, rheumatism, expectorant, and febrifuge properties (http://www.naturalmedicinalherbs.net/herbs/natural/).

## 5. Conclusion

In this manuscript, we have provided an *in silico* evidence for the ability of miRNAs to target *S. aureus* virulence and resistance genes. The predicted miRNAs have also been verified for target site accessibility, suggesting their higher likelihood to target *S. aureus* transcriptome. Further investigations on the intracellular stability of these miRNAs and their purported anti-resistance and anti-virulence activity under in vitro and in vivo conditions have to be performed. In the future, comprehensive experimental validation of these issues will aid in the development of efficient and non-toxic therapy for *S. aureus*.

