## Supplementary material for "Plant miRNAs Anti-*Staphylococcus aureus:* Therapeutic Perspective": Table S2

| **miRNAs** | **Minimum free energy (MFE)** | **MFE secondary structure** |
| --- | --- | --- |
| >gma-miR1524 MIMAT0007385 Glycine max miR1524  CGAGUCCGAGGAAGGAACUCC | **-7.03** kcal/mol  (stable) | 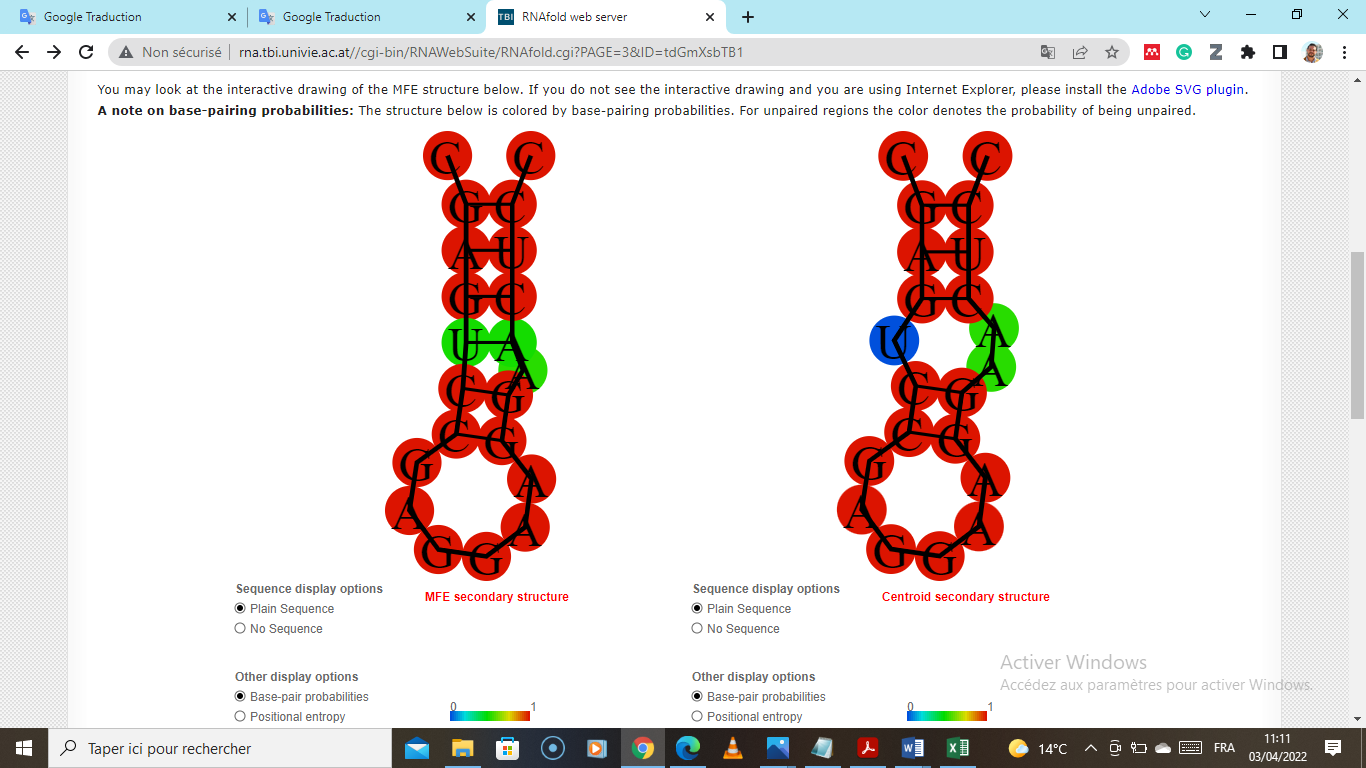 |
| >ppt-miR1074 MIMAT0005204 Physcomitrella patens miR1074  AGGGUUGUUAGUUGUGUUGAU | **-0.13** kcal/mol (unstable) | 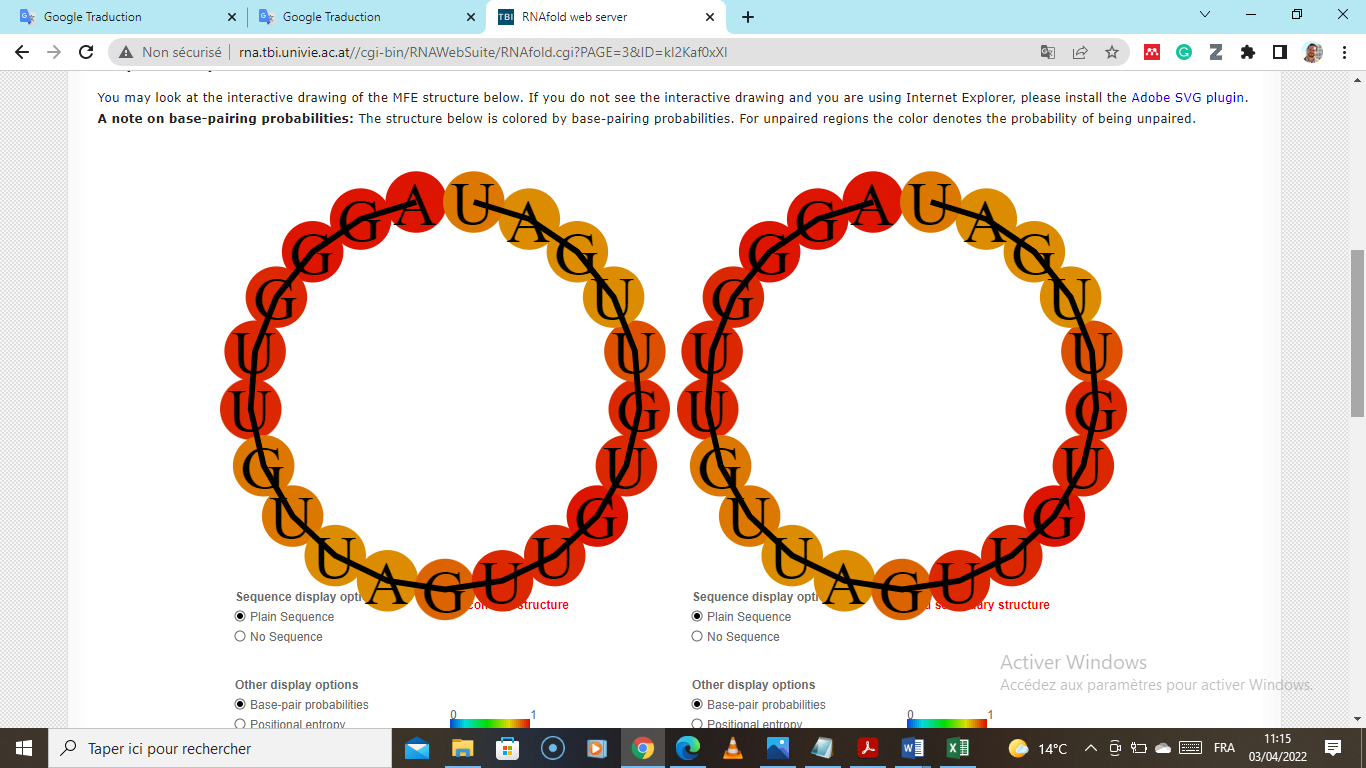 |
| >ath-miR8181 MIMAT0032780 Arabidopsis thaliana miR8181  UGGGGGUGGGGGGGUGACAG | **-0.16** kcal/mol (unstable) | 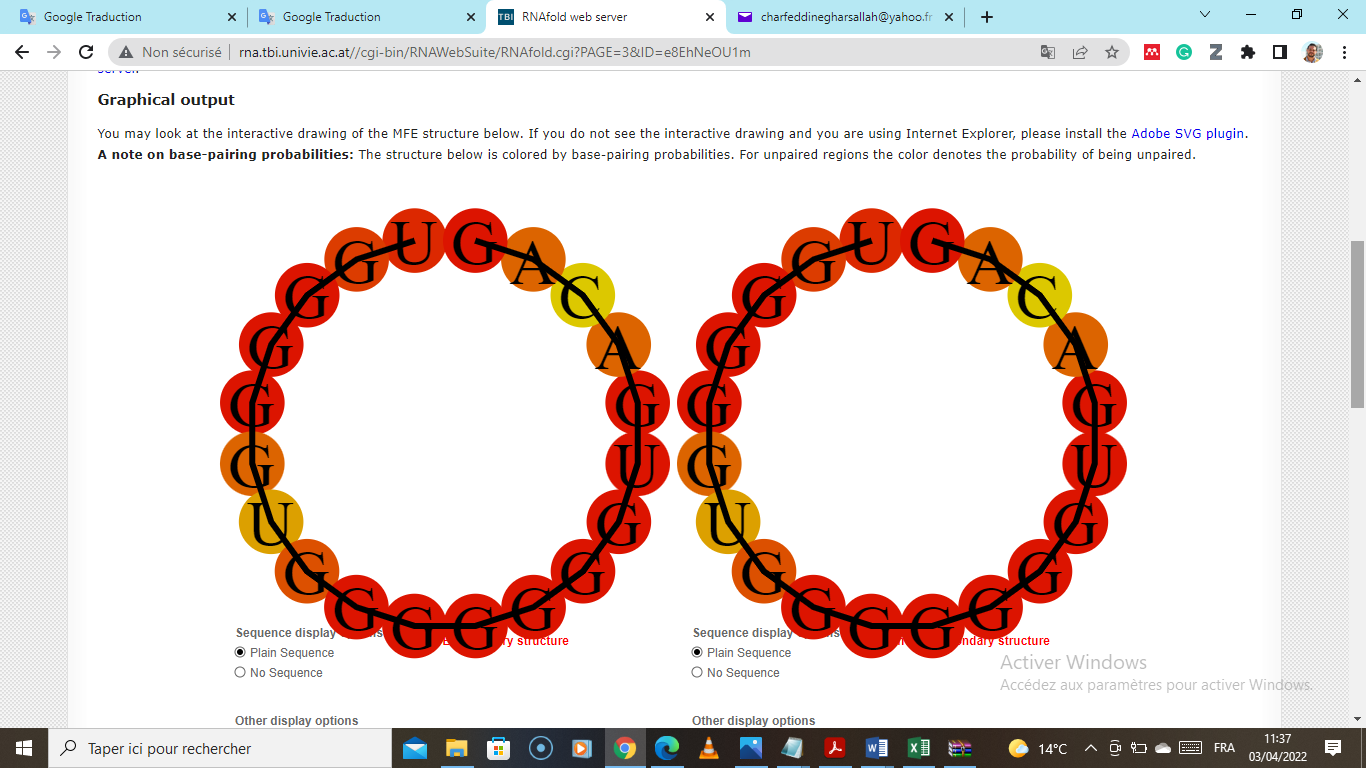 |
| >csi-miR502a-3p  GGGUUUUGAGUGGAAACUGAGU | **-3.28** kcal/mol  (stable) | 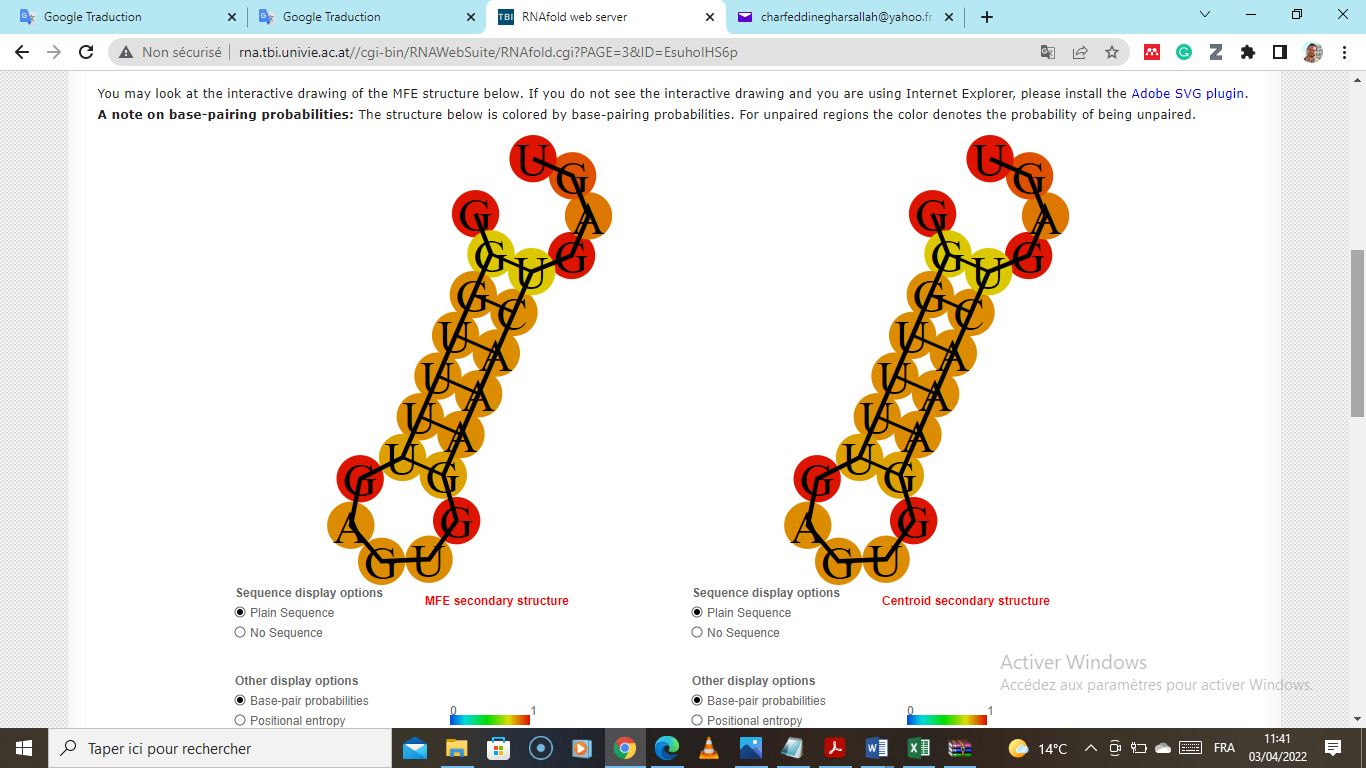 |
| >ath-miR775 MIMAT0003934 Arabidopsis thaliana miR775  UUCGAUGUCUAGCAGUGCCA | **-0.36** kcal/mol (unstable) | 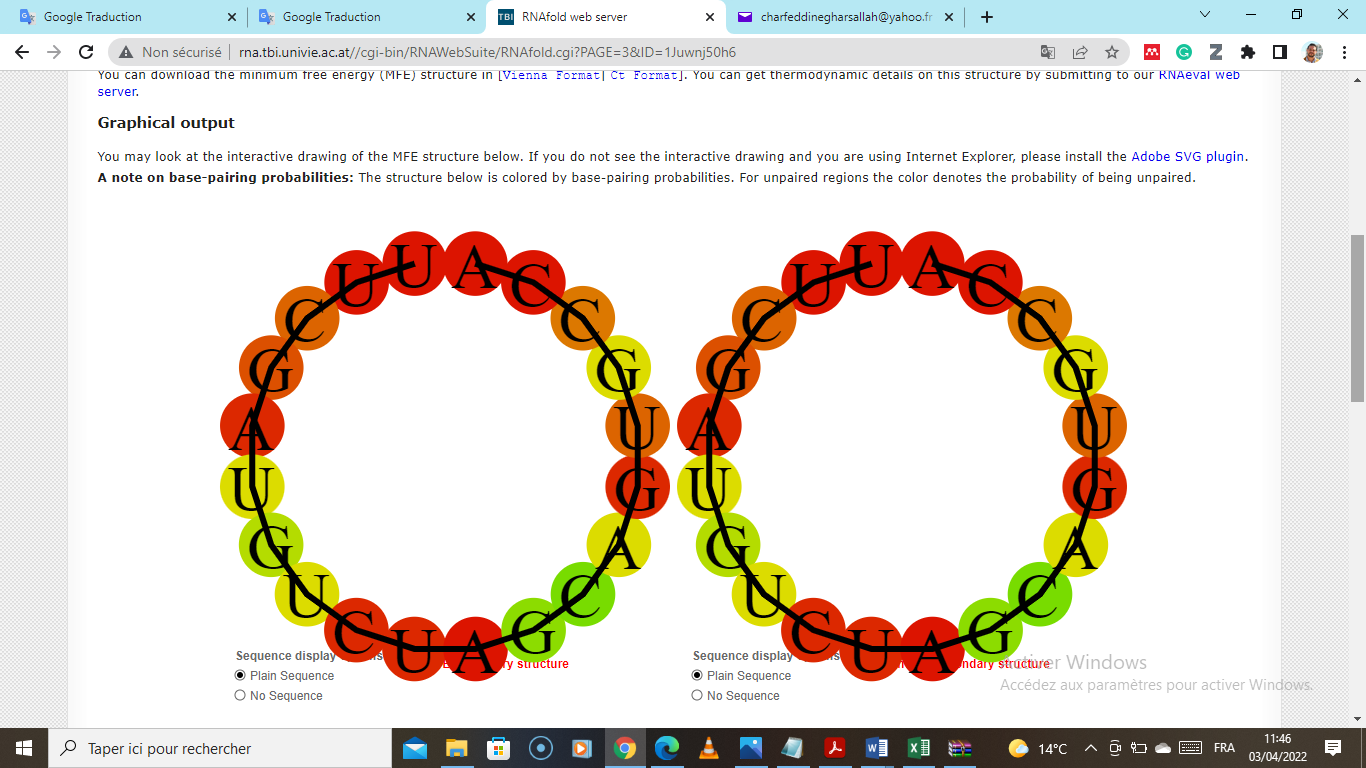 |
| >csi-miR451-5p  AGGCGGUCGUCUUUUGGCUAAC | **-3.45** kcal/mol  (stable) | 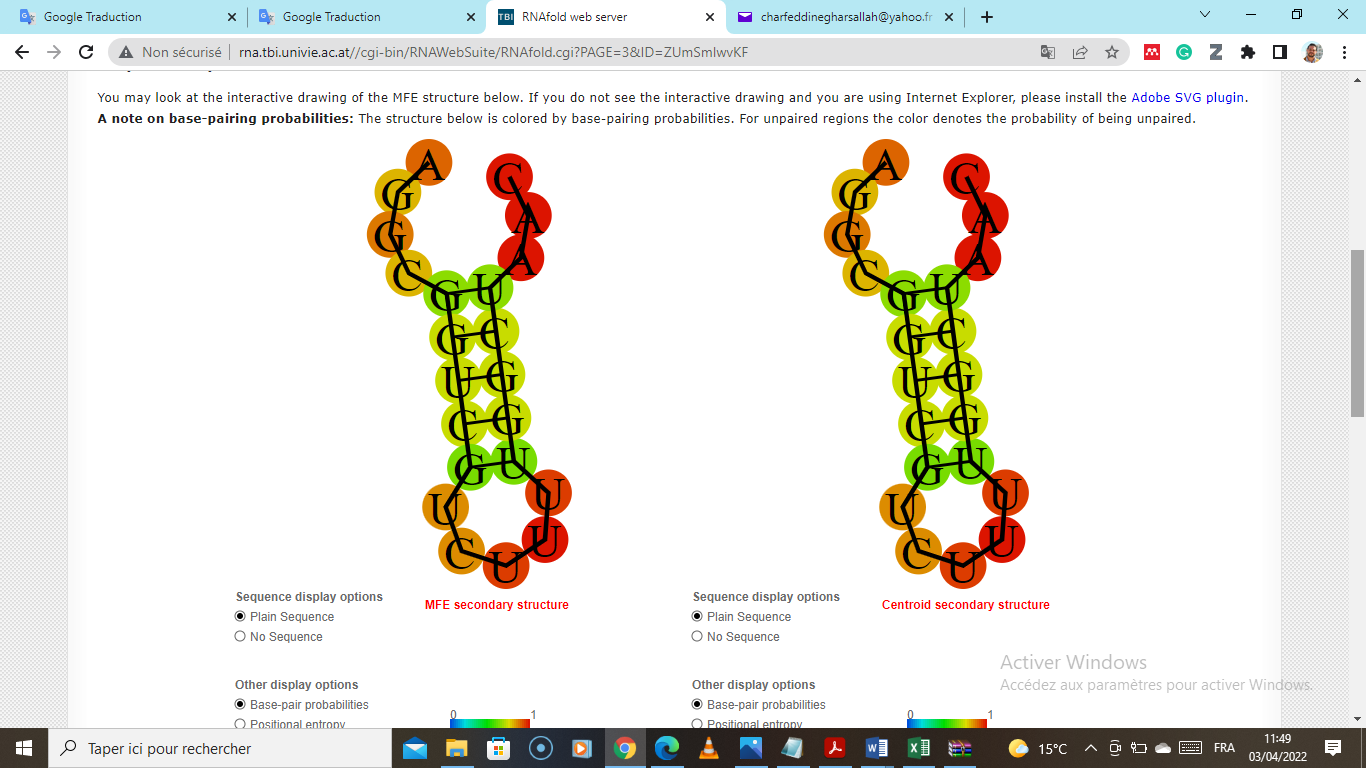 |
| >aly-miR166f-3p MIMAT0017470 Arabidopsis lyrata miR166f-3p  UCGGACCAGGCUUCAUUCCCC | **-0.93** kcal/mol  (stable) | 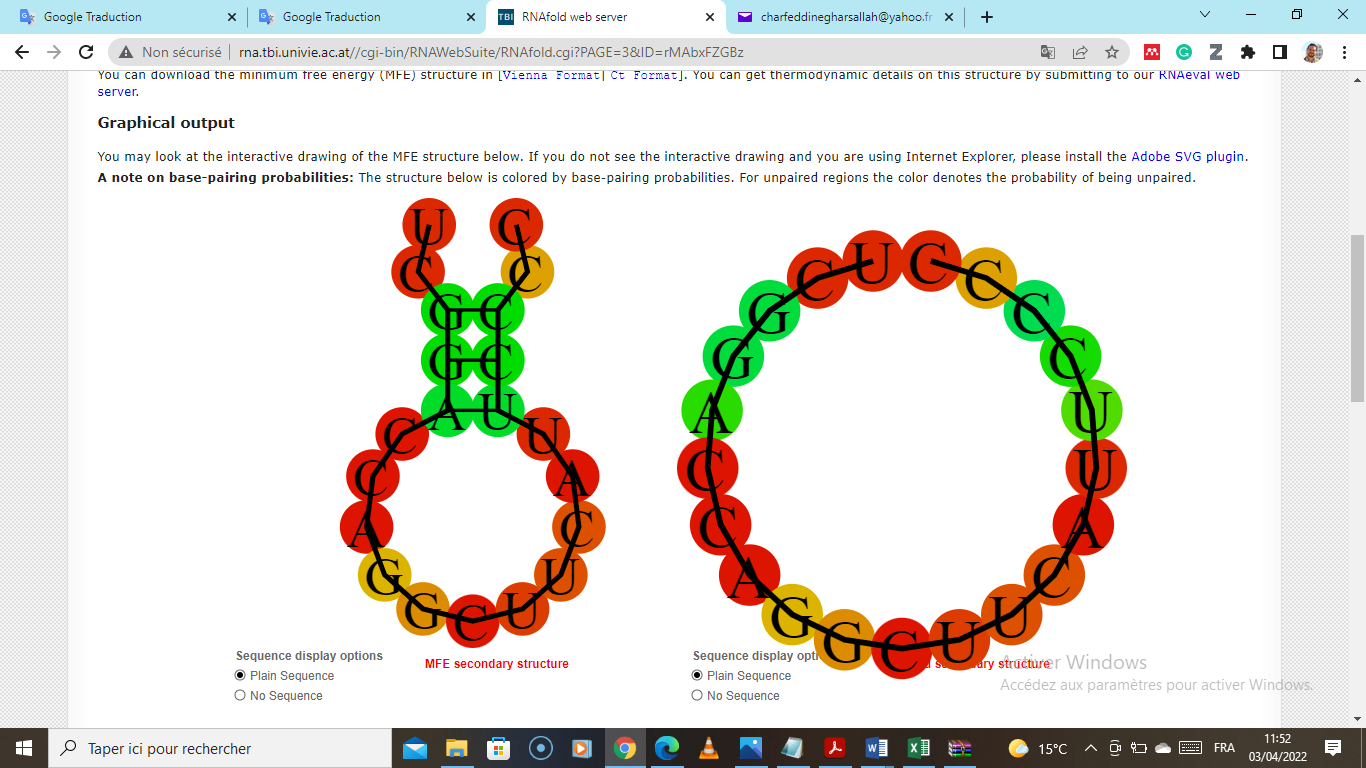 |
| >csi-miR3953 MIMAT0018496 Citrus sinensis miR3953  UUGAGUUCUGCAAGCCGUCGA | **-0.00** kcal/mol  (unstable) | 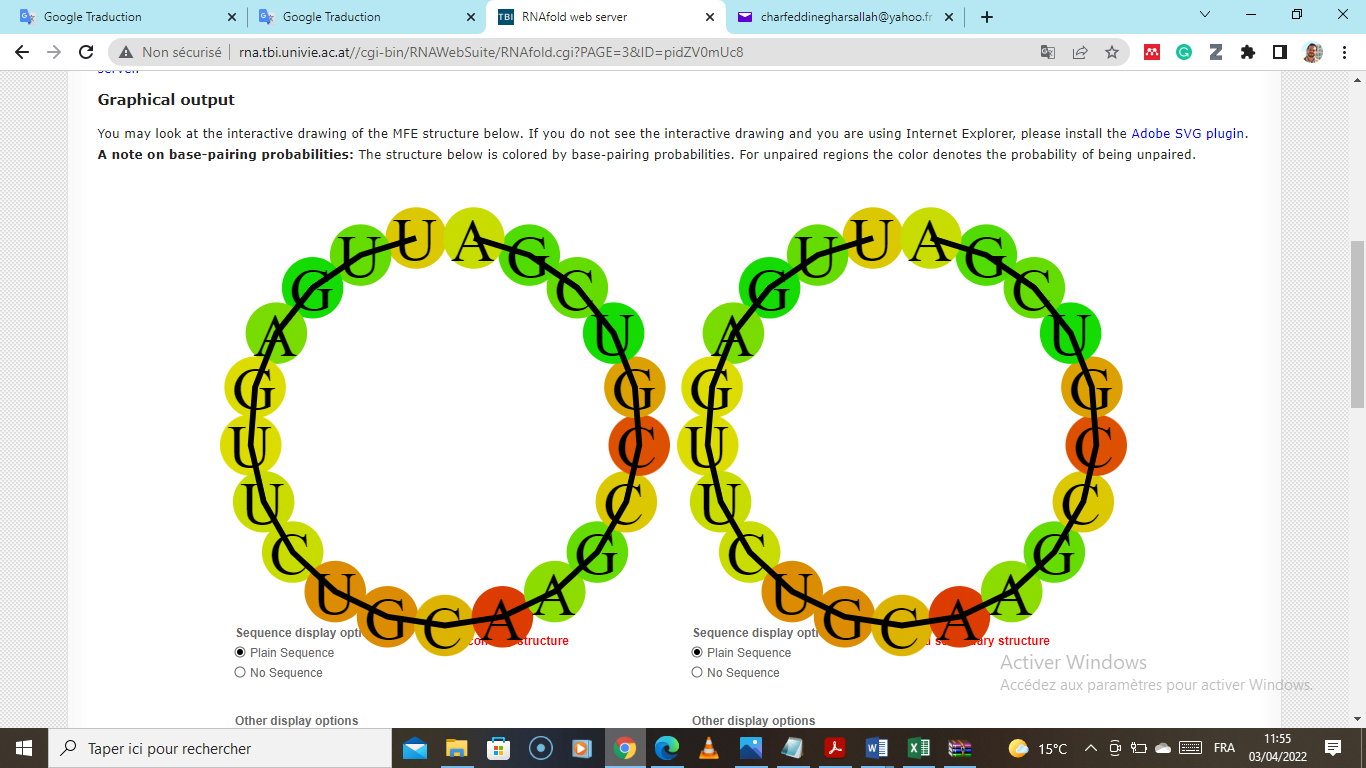 |
| >csi-miR263-3p  AUGUAUGGAUGGAUGGAUGGAUG | **-0.72** kcal/mol  (stable) | 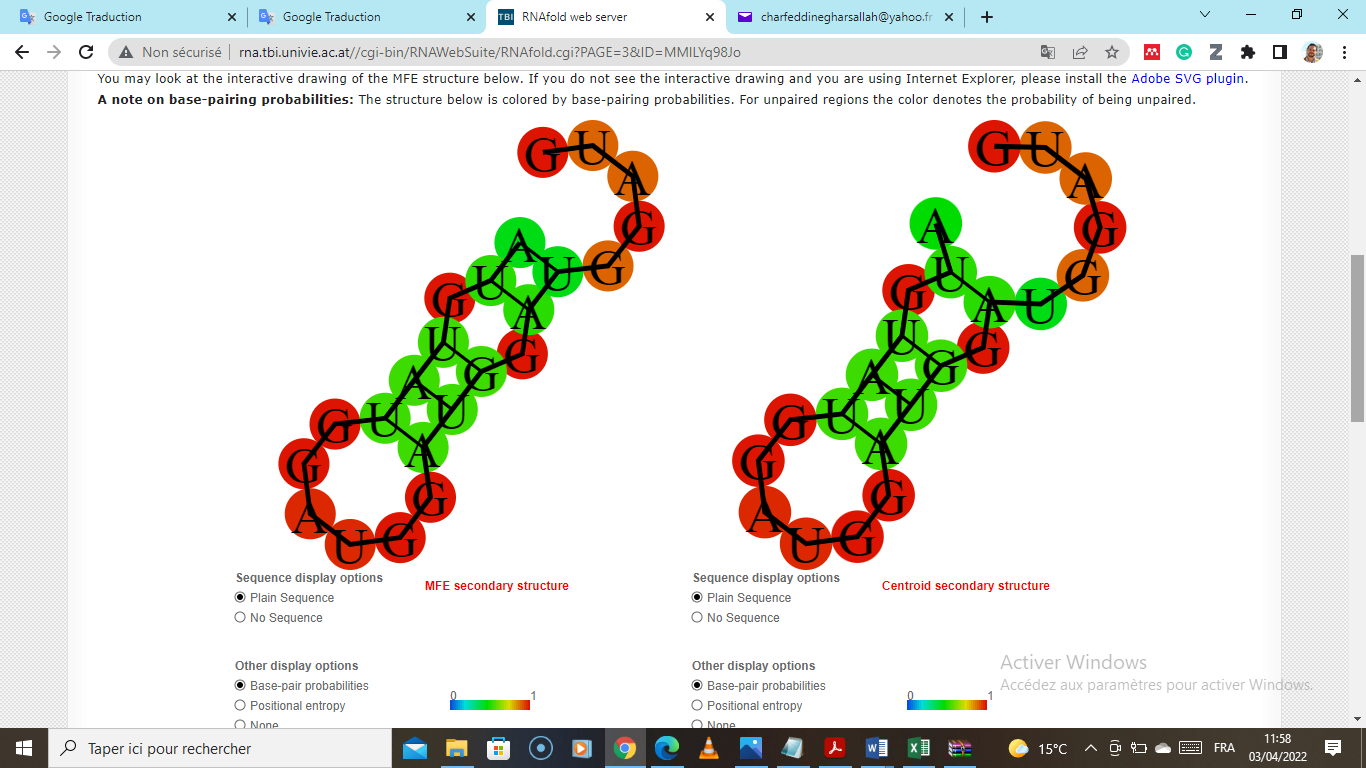 |
| >csi-miR807  UUGAGUUCUGCAAGCCGUCGA | **-0.00** kcal/mol  (unstable) | 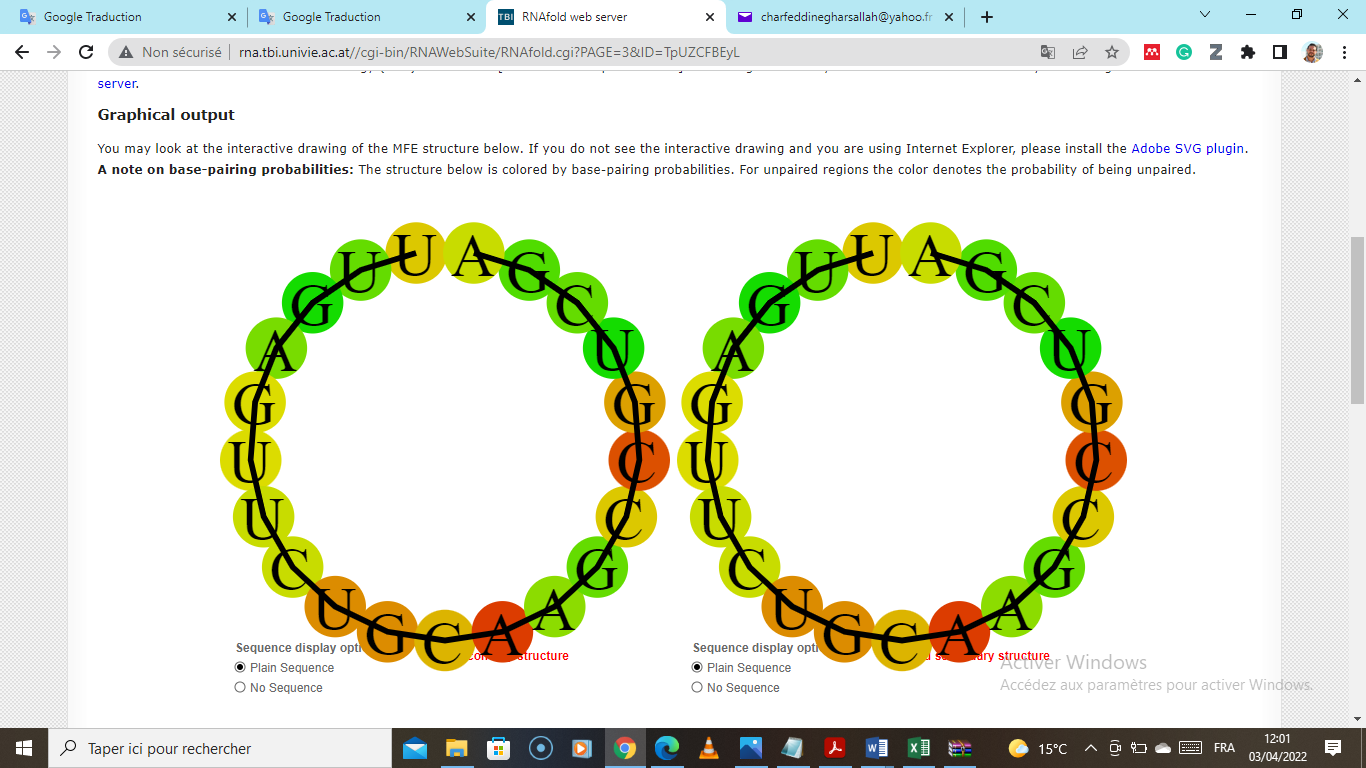 |
| >mtr-miR5554a-3p MIMAT0022200 Medicago truncatula miR5554a-3p  ACCAUCGUUGCAGAUGCUCAUC | **-3.23** kcal/mol  (stable) | 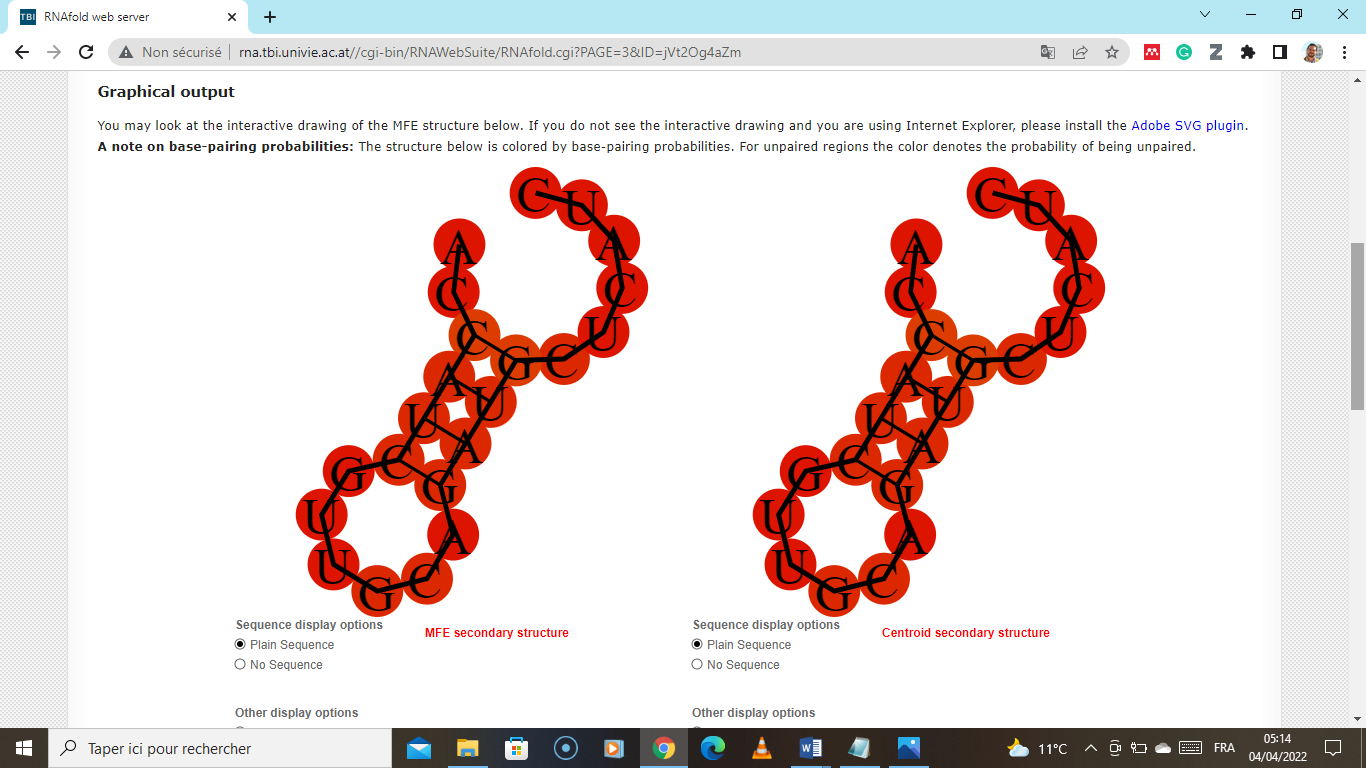 |
| >mtr-miR5244 MIMAT0021240 Medicago truncatula miR5244  UAUCUCAUGAAGAUUGUUGGU | **-0.80** kcal/mol  (stable) | 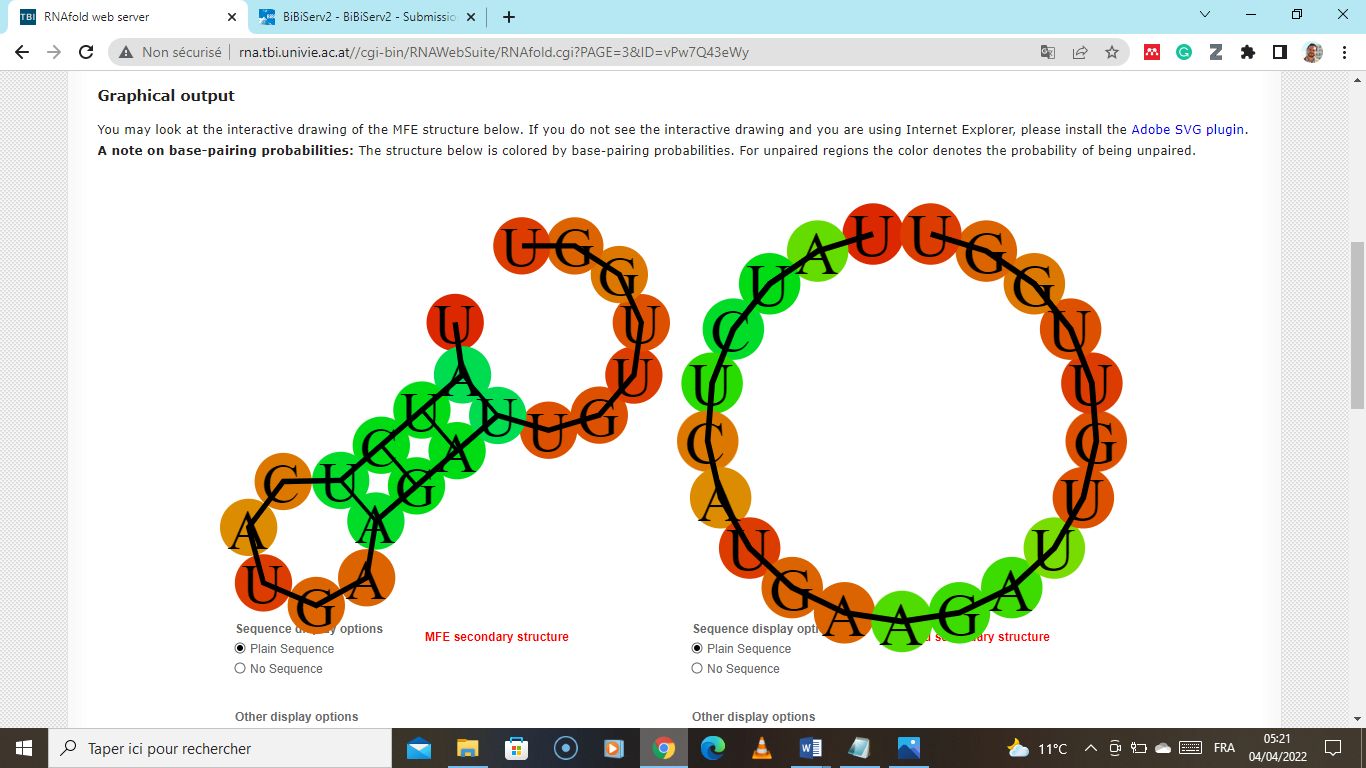 |
| >mtr-miR399t-5p MIMAT0030005 Medicago truncatula miR399t-5p  GGGUGAGUUCUCCAUUGGCAGGU | **-1.00** kcal/mol  (unstable) | 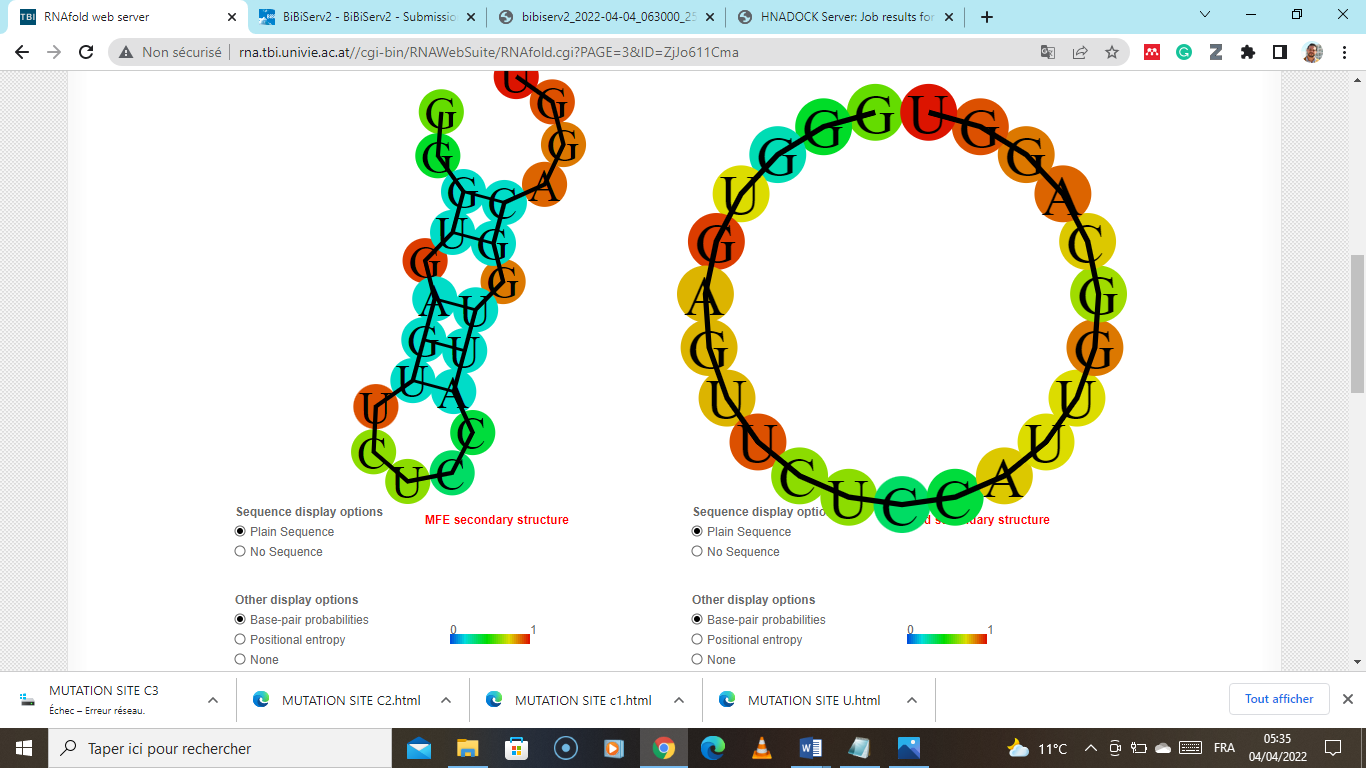 |
| >osa-miR6256 MIMAT0024879 Oryza sativa miR6256  GUAGUACUCGGUUGUAGGUGUA | **-0.50** kcal/mol  (unstable) | 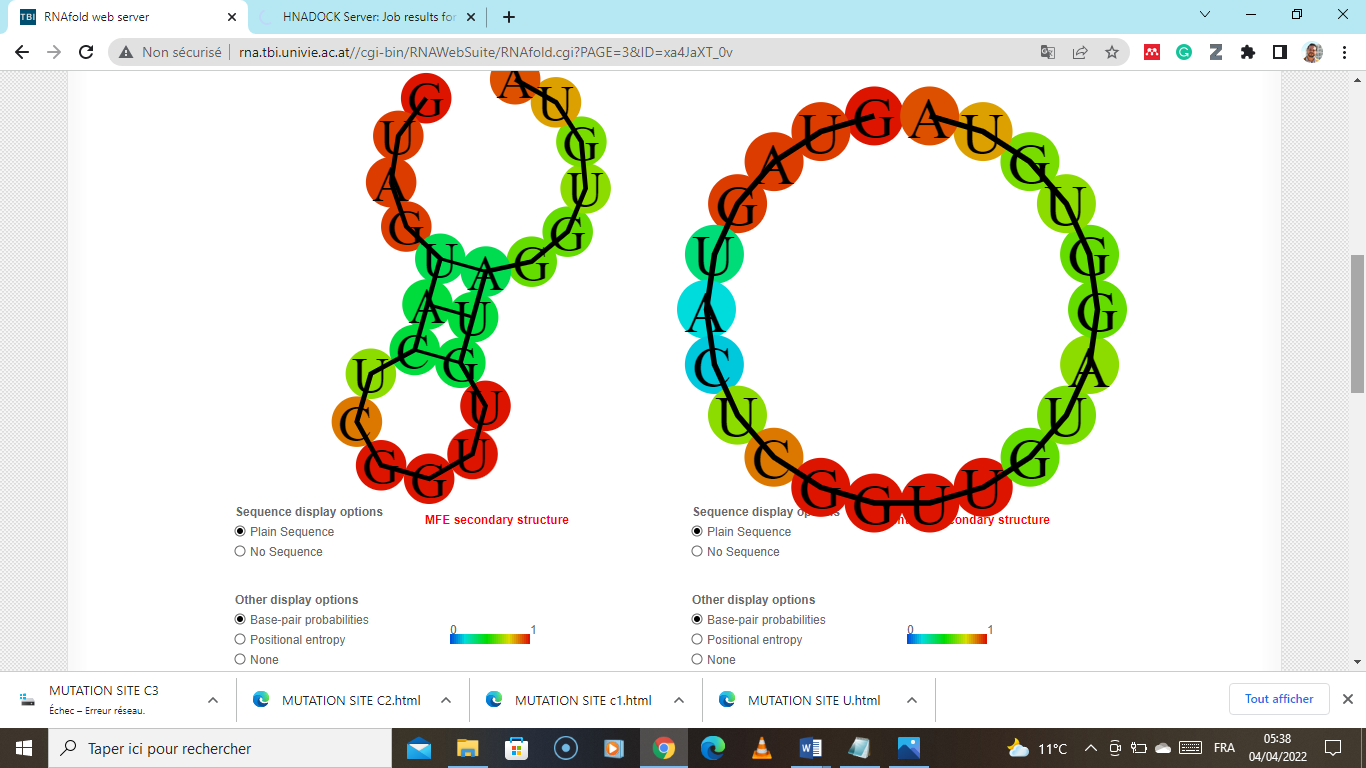 |
| >ptc-miR171g-5p MIMAT0022902 Populus trichocarpa miR171g-5p  UGUUGGGAUGGCUCAAUCAUG | **-4.95** kcal/mol  (stable) | 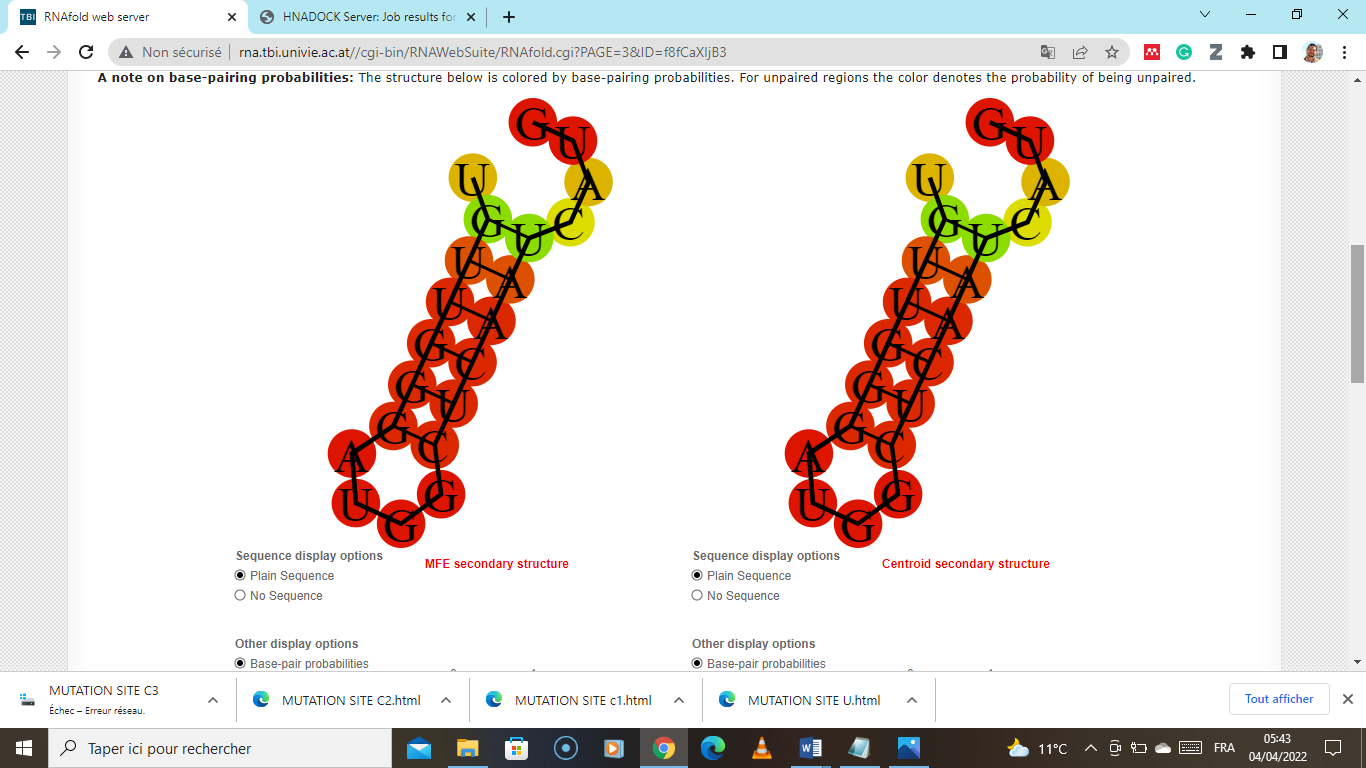 |
| >ptc-miR171h-5p MIMAT0022903 Populus trichocarpa miR171h-5p  UGUUGGGAUGGCUCAAUCAUA | **-4.95** kcal/mol  (stable) | 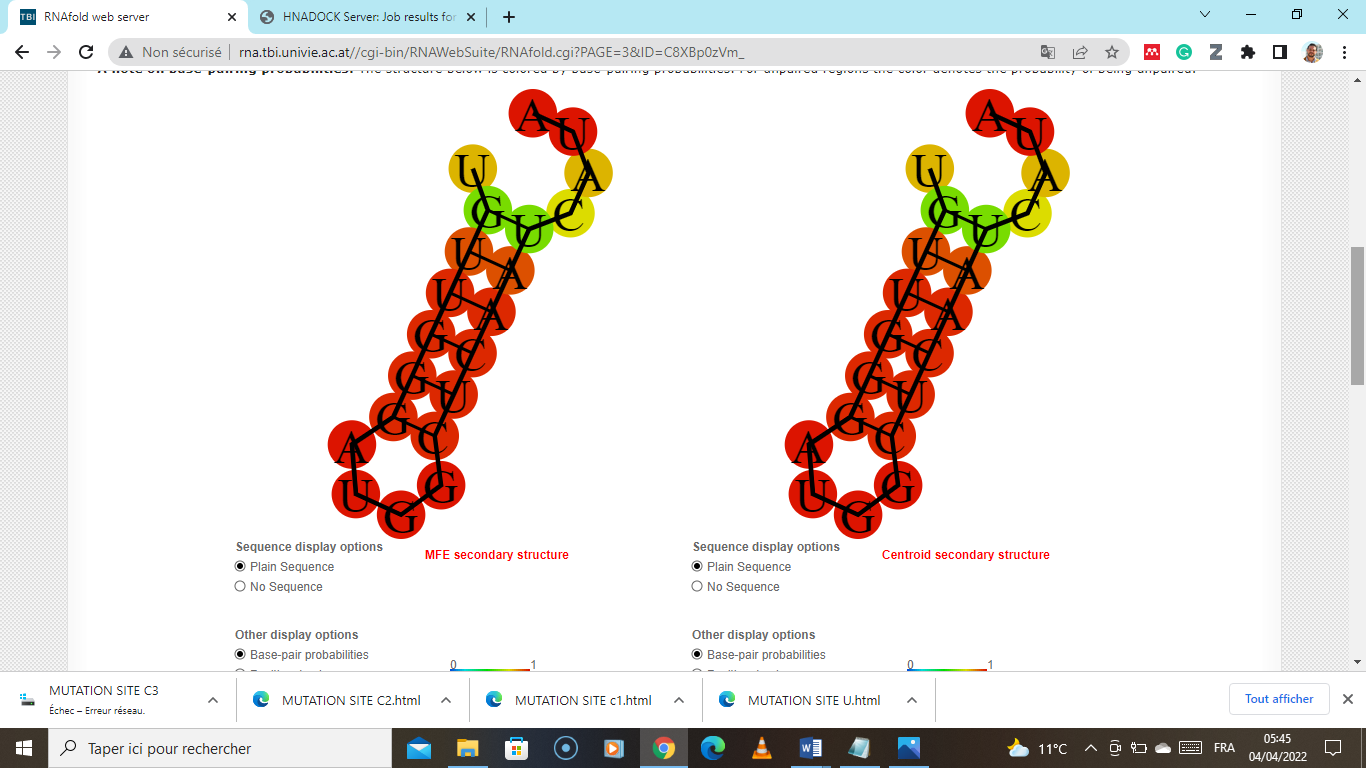 |
| >stu-miR8045 MIMAT0030959 Solanum tuberosum miR8045  AUUGAUAGUUGAGGUGUGUUU | **-0.13** kcal/mol  (unstable) | 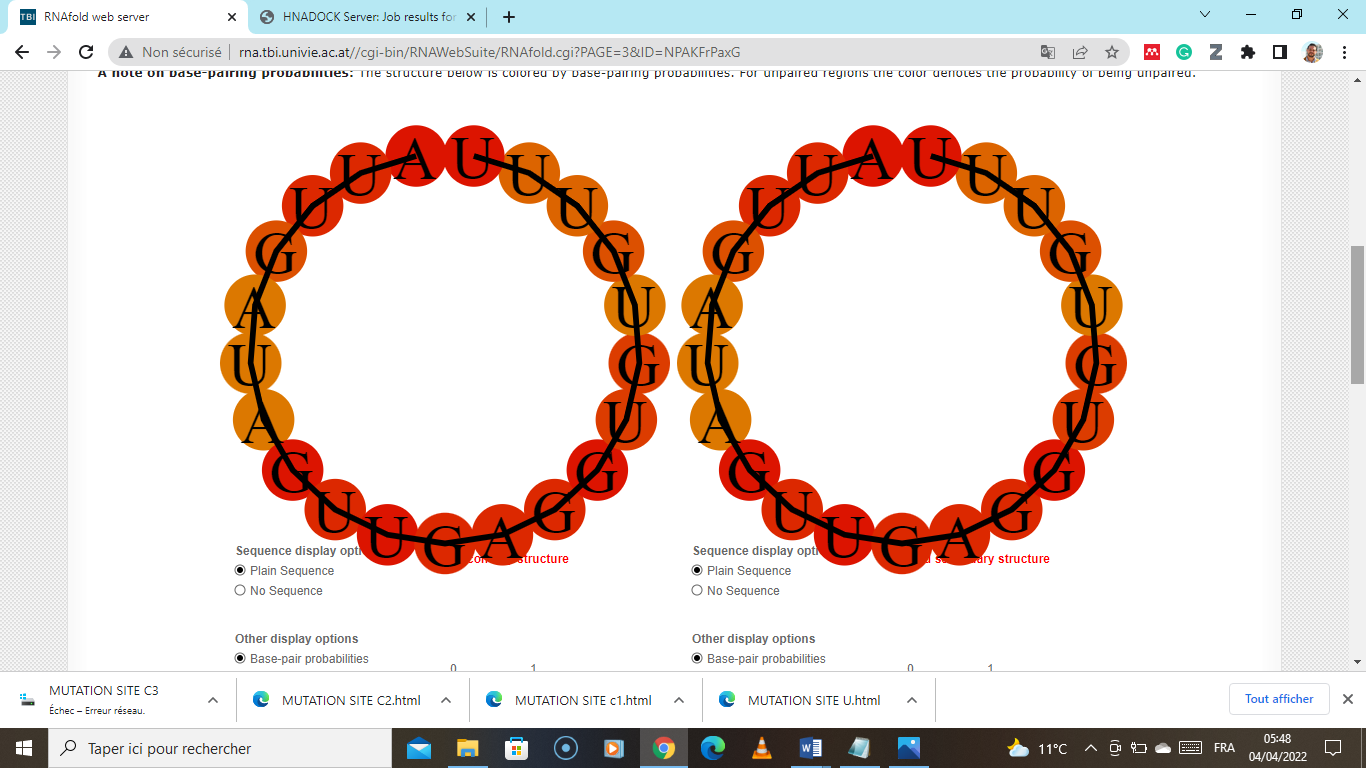 |
| >smo-miR1097 MIMAT0005253 Selaginella moellendorffii miR1097  UAGCCAUUGUUGUUGUUGGAA | **-1.89** kcal/mol  (stable) | 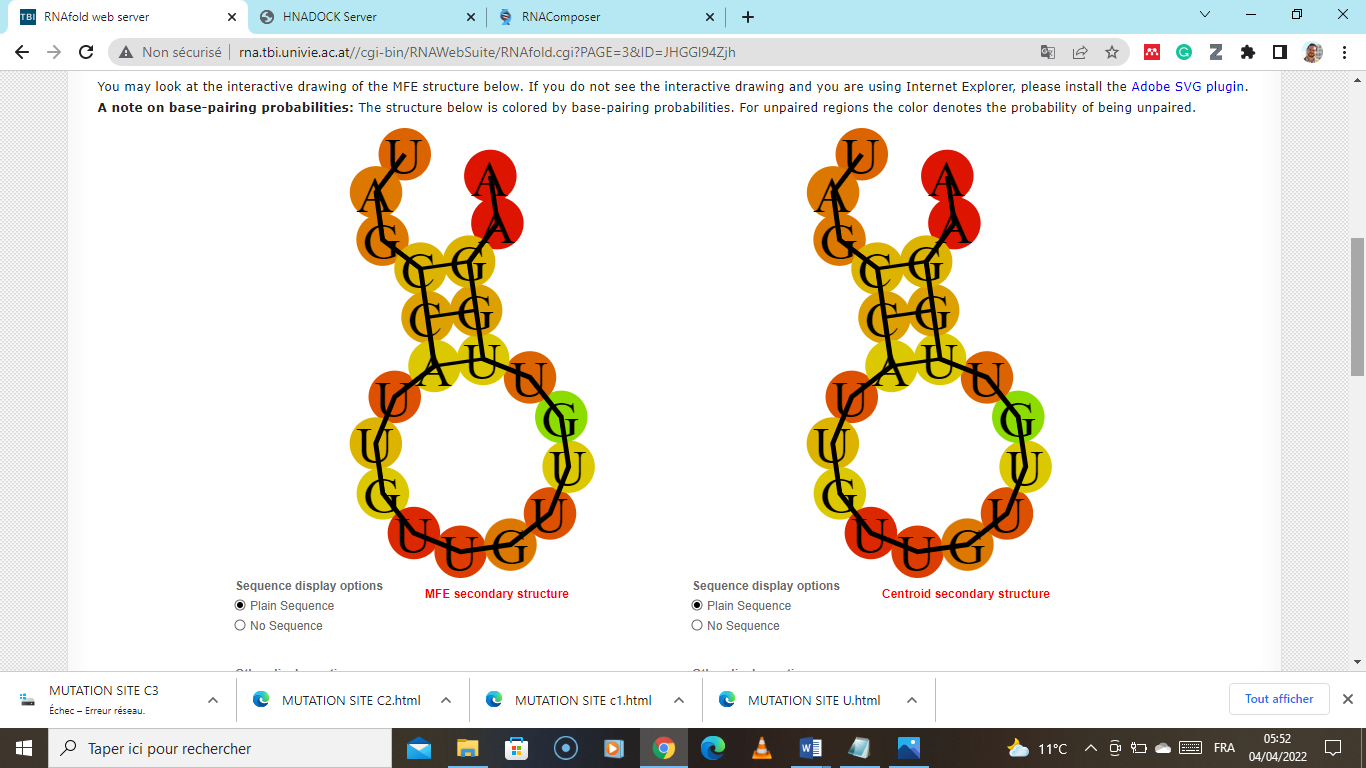 |
| >smo-miR1093 MIMAT0005246 Selaginella moellendorffii miR1093  UGGAGGUGUCGUUGCCAAGGA | **-2.82** kcal/mol  (stable) | 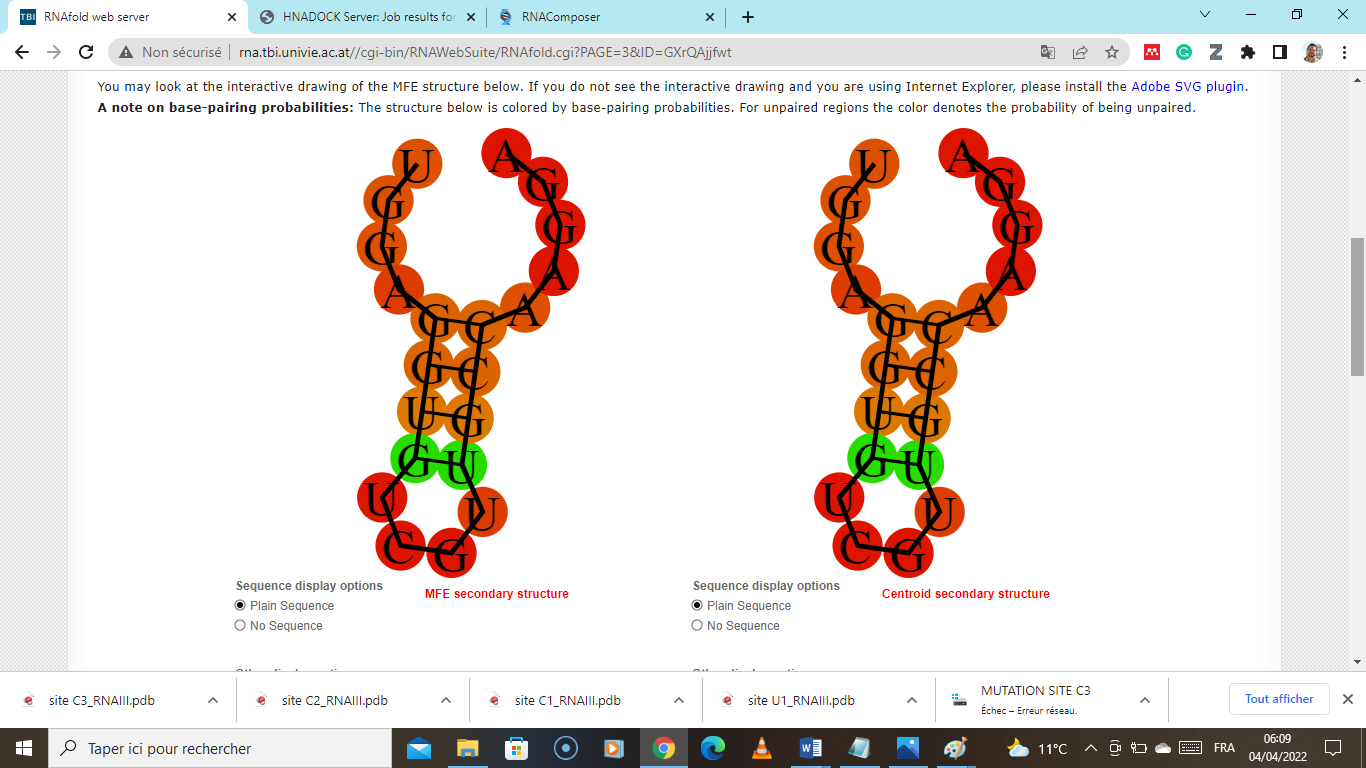 |
| >bna-miR6032 MIMAT0023652 Brassica napus miR6032  UGGAGCAUCAACAGAUCUCGG | **-2.47** kcal/mol  (stable) | 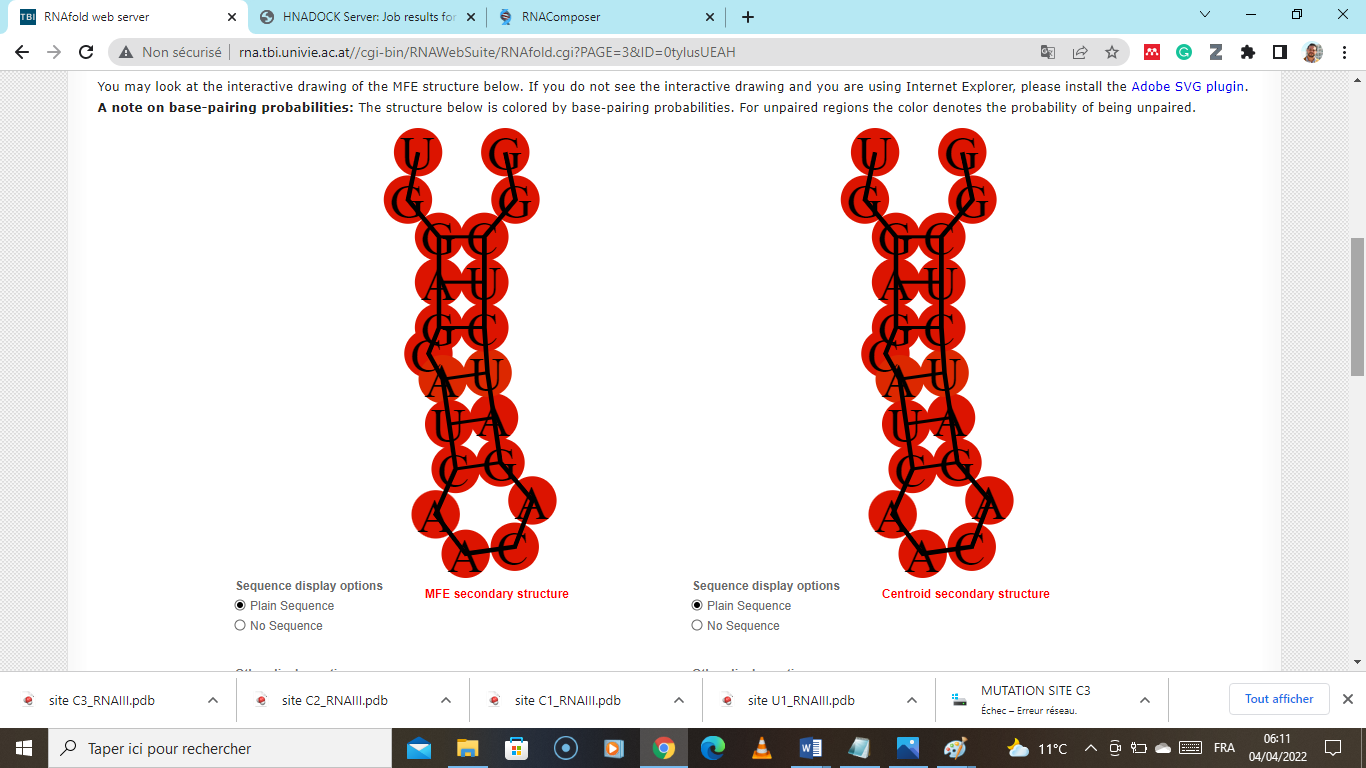 |
| >amo-miR16-5p  UUUGUGCCAUUGAUCUGAUAAA | 0.0 kcal/mol   (unstable) | 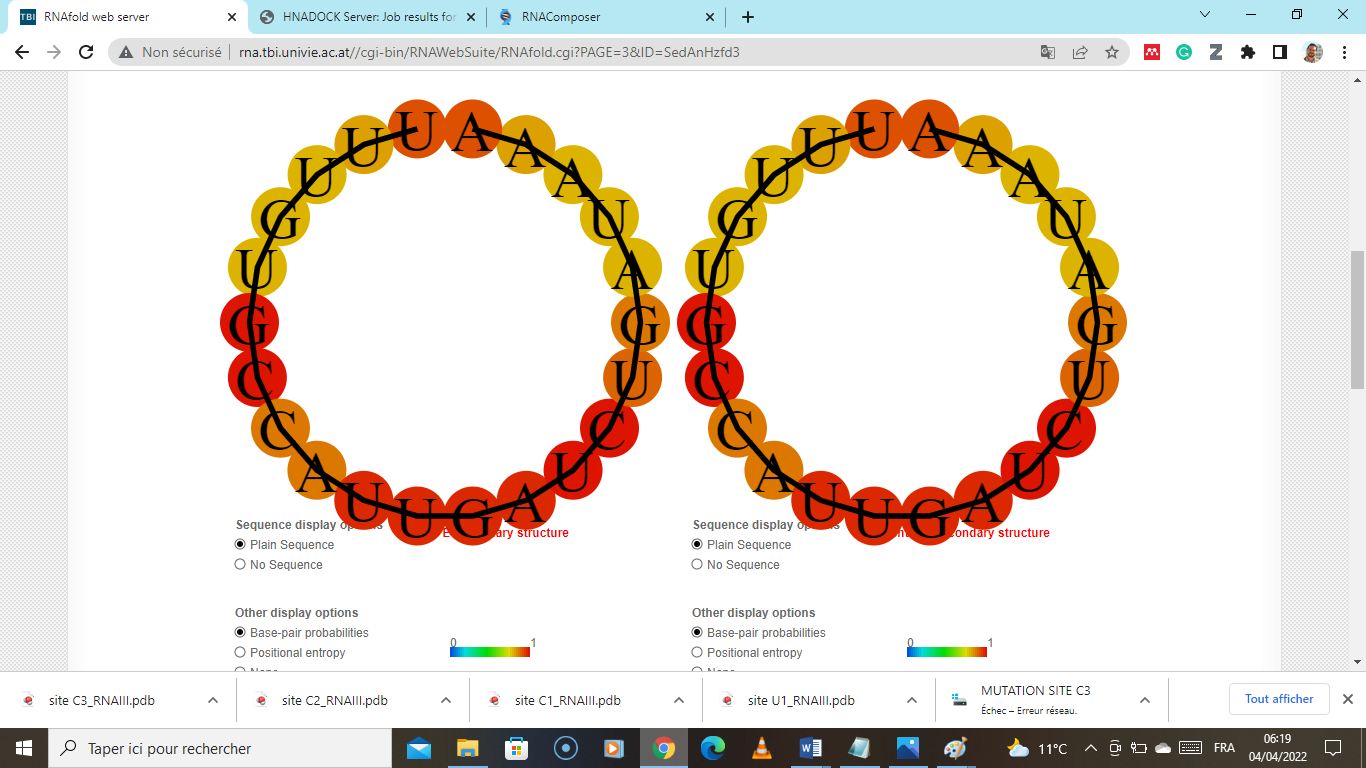 |
| >amo-miR113-5p  CCUUCCAACUAUGUGUCUUU | 0.0 kcal/mol   (unstable) | 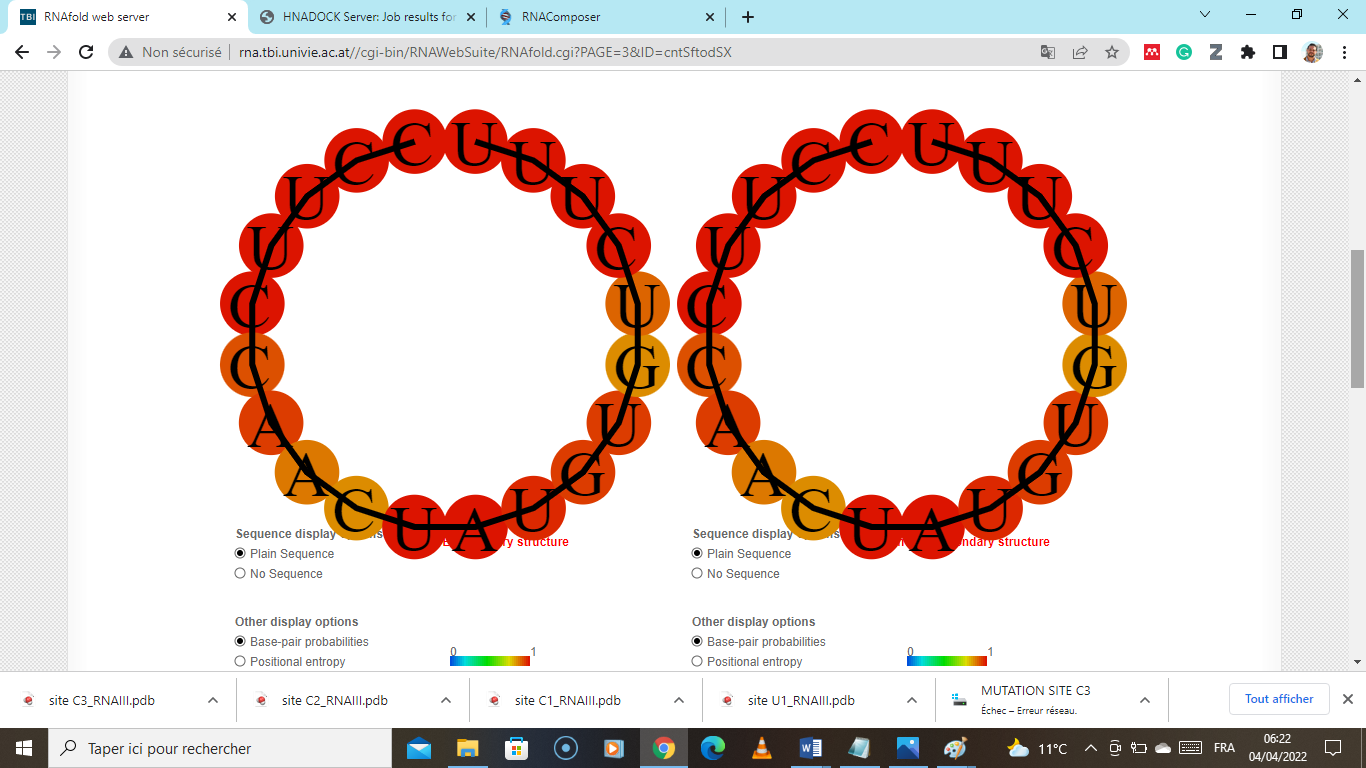 |
| >gbi-miR373  AUUGAUAGUUGAGGUGUGUUU | 0.0 kcal/mol   (unstable) | 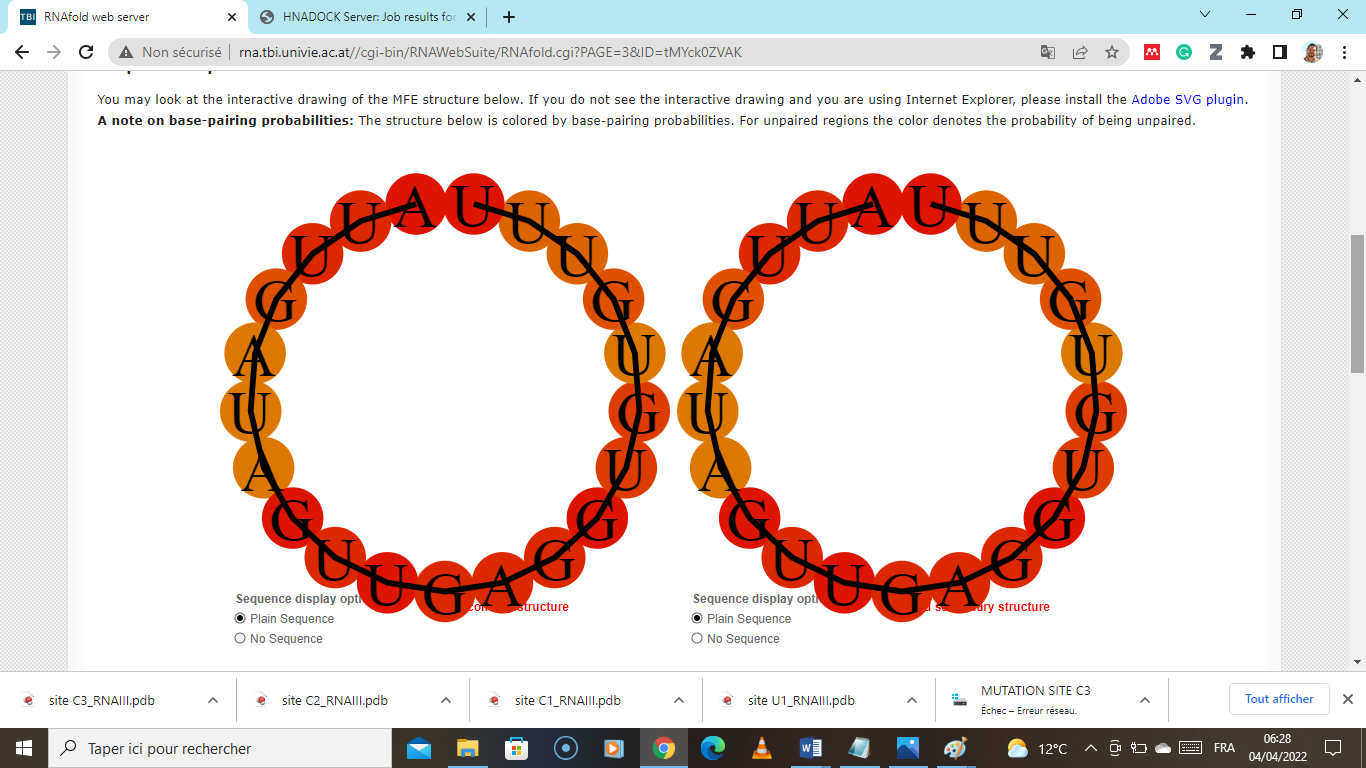 |
| >dmo-miR40-5p  CUGAUGUUGAAAUCGCCUCUCC | **-1.66** kcal/mol  (stable) | 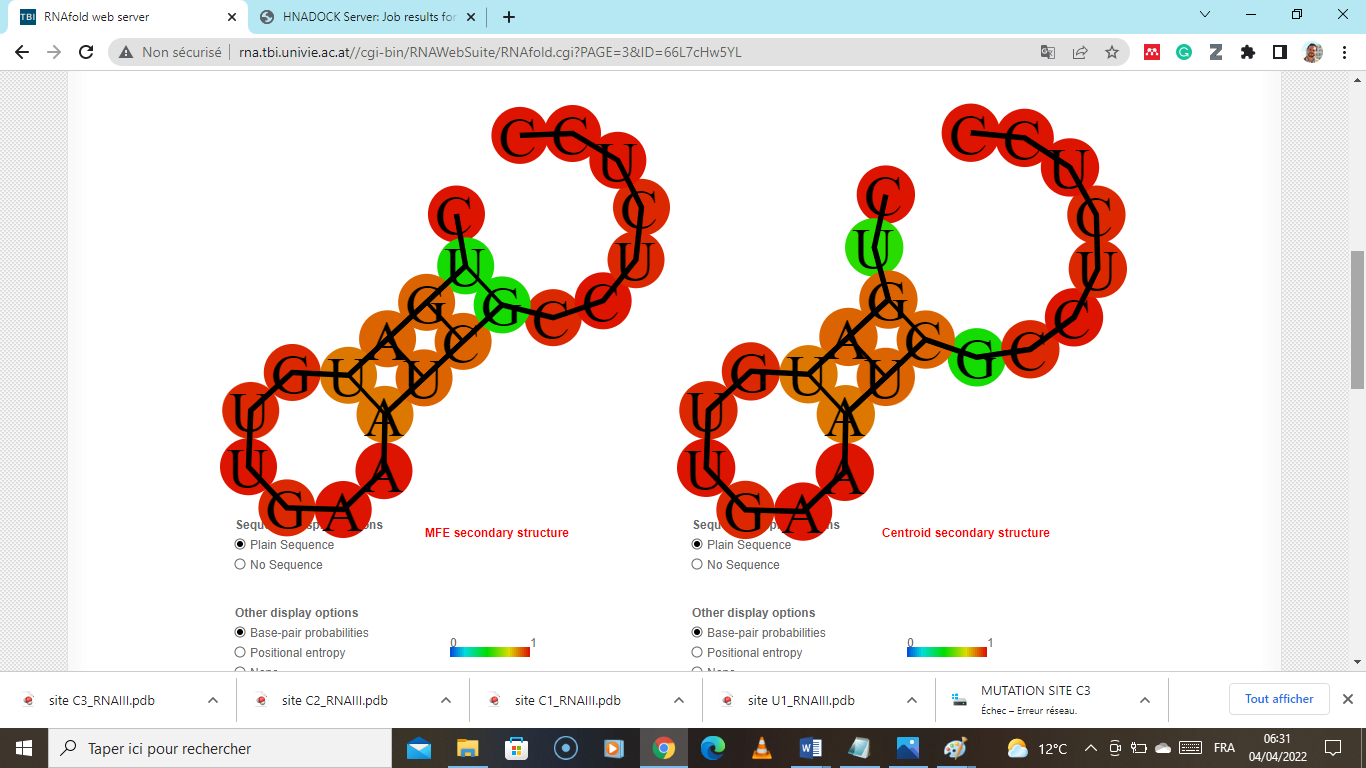 |
| >dof-miR12-3p  UUGGGAUGGCUUUUUUACUGC | 0.0 kcal/mol   (unstable) | 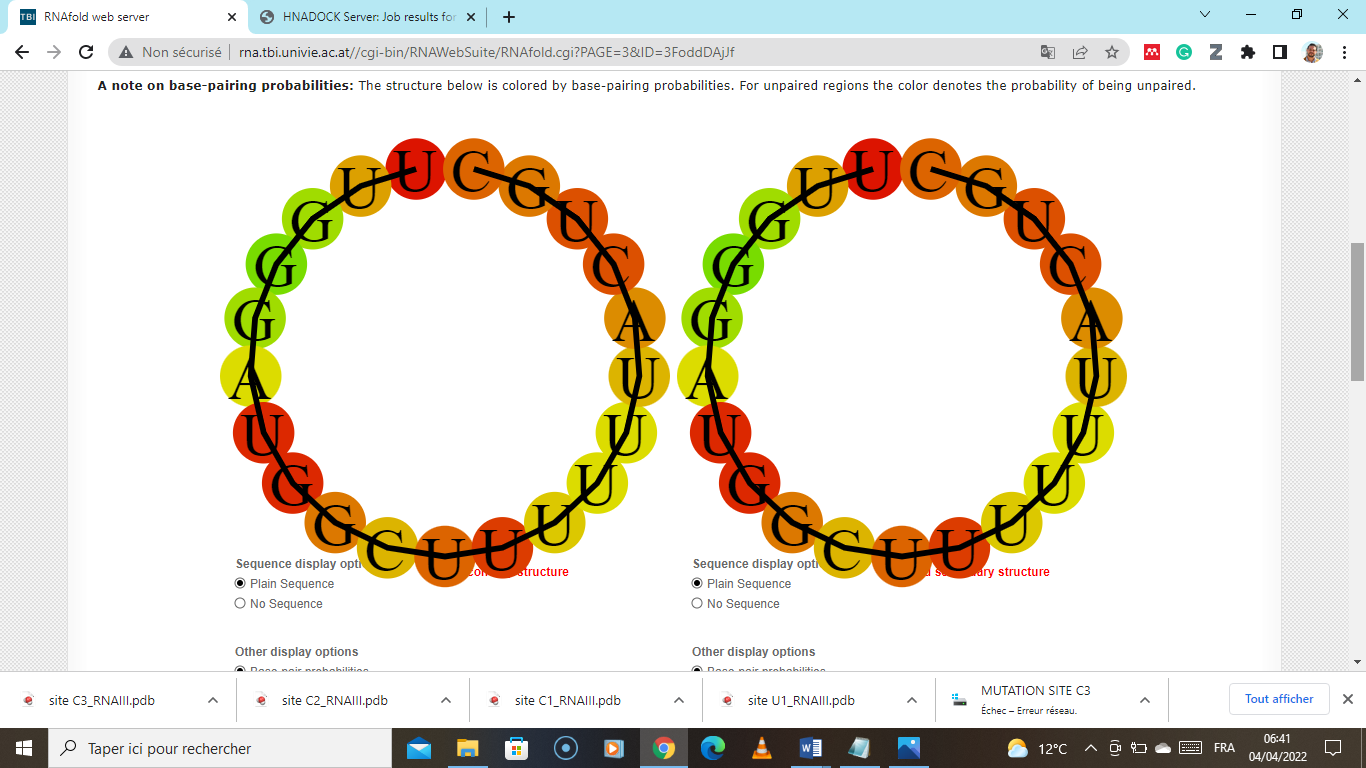 |
| >lyc-miR30-3p  CGGAGGAUGGUUAUGUUCACCG | **-4.63** kcal/mol  (stable) | 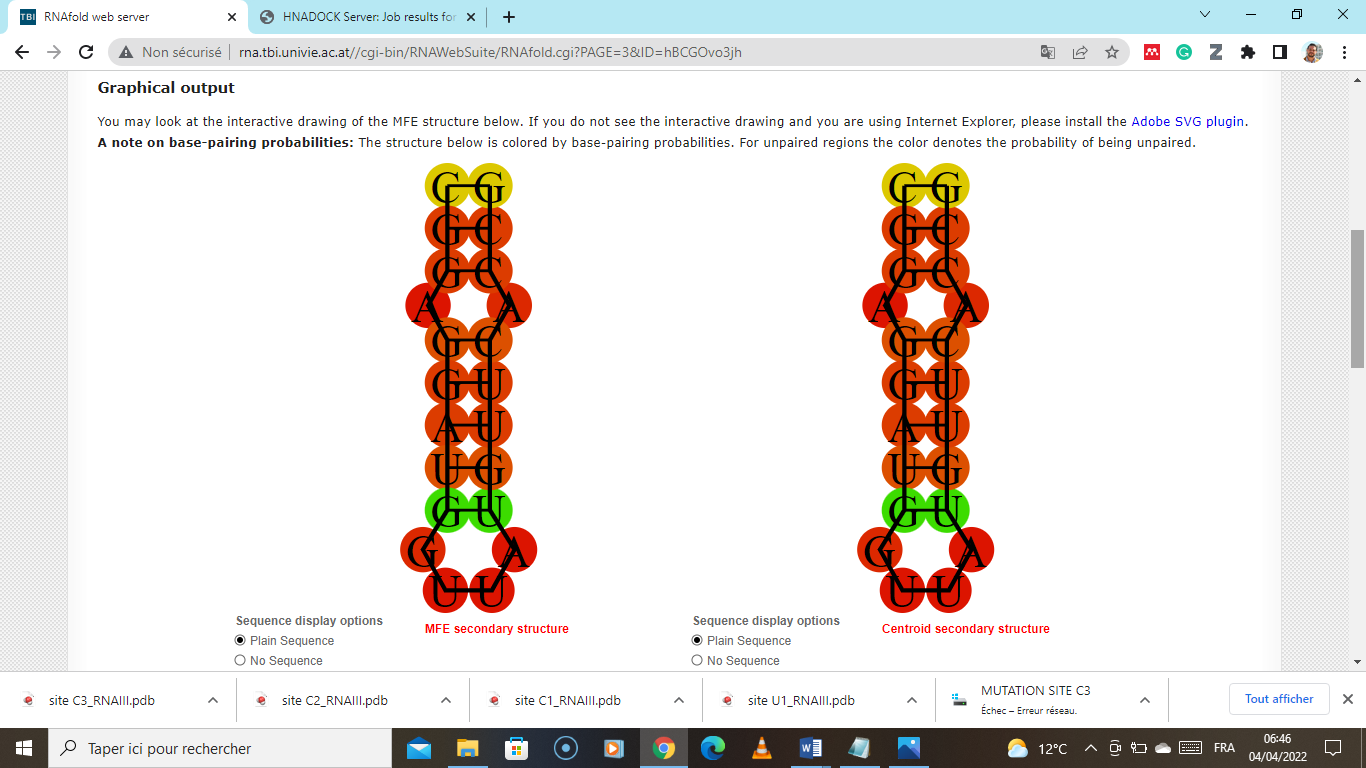 |
| >smi-miR112-3p  CAGCUUGUGGAAGCAUCUGA | **-4.02** kcal/mol  (stable) | 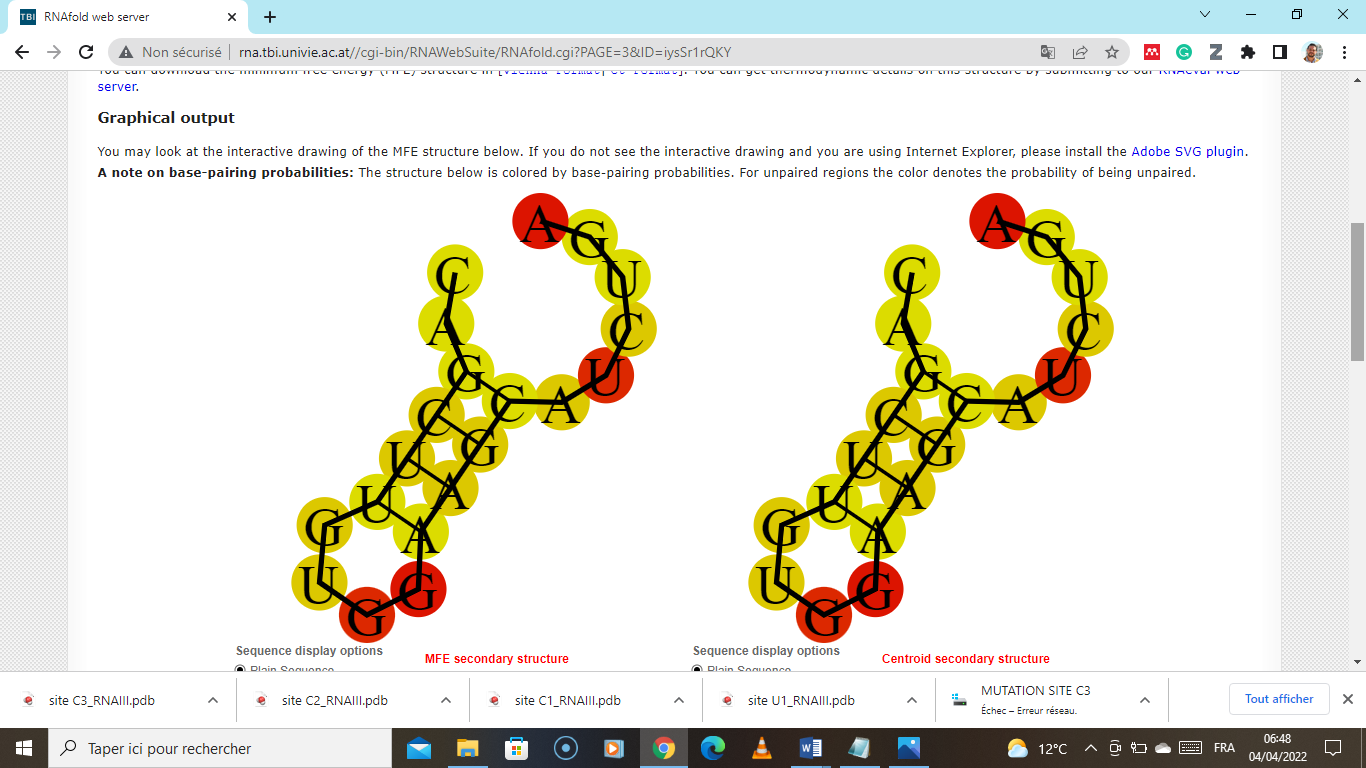 |
| >pgr-miR2-3p  CCAACUUCUCUACAUCCUCUU | **-0.00** kcal/mol  (unstable) | 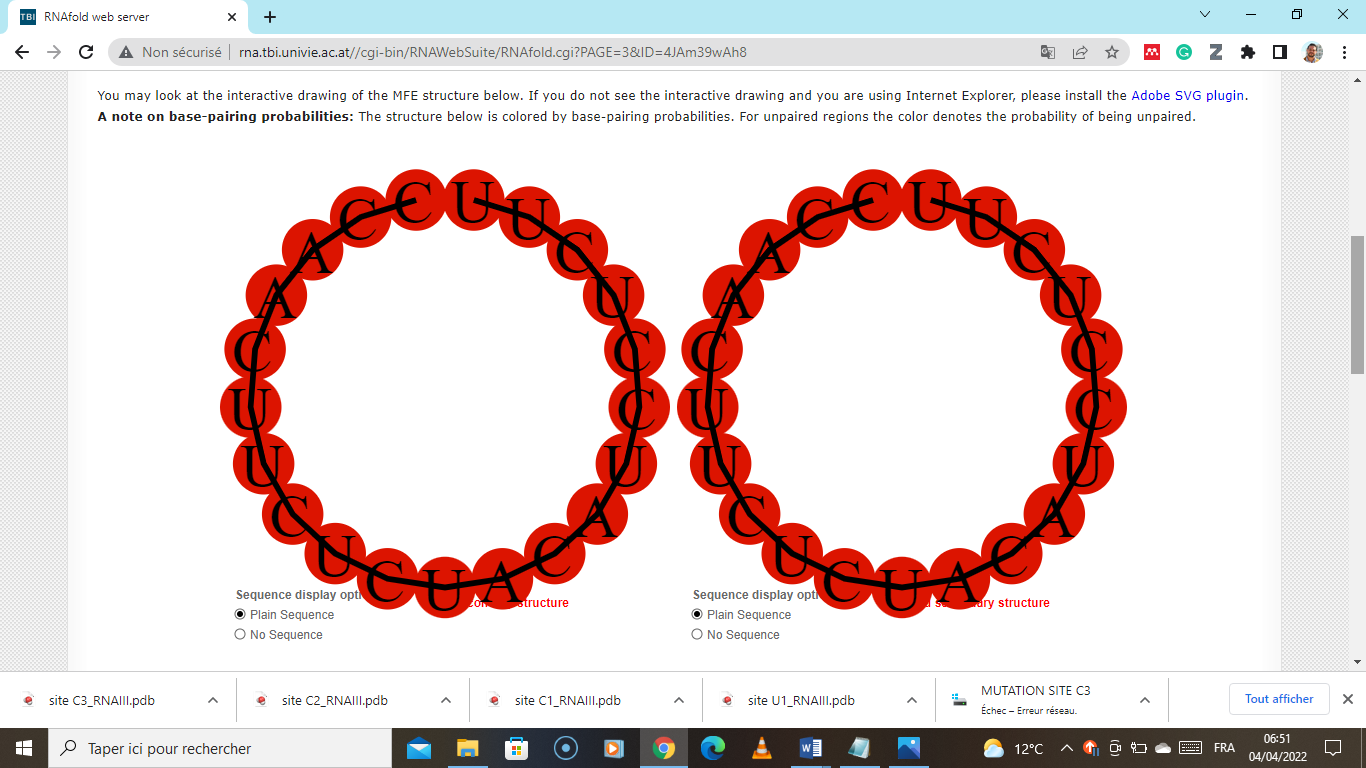 |

### Table S2: Results for thermodynamic ensemble prediction
